# Integrative multi-omics approach for stratification of tumor recurrence risk groups of Hepatocellular Carcinoma patients

**DOI:** 10.1101/2021.03.03.433841

**Authors:** Harpreet Kaur, Anjali Lathwal, Gajendra P.S. Raghava

## Abstract

Postoperative tumor recurrence is one of the major concerns associated with the poor prognosis of HCC patients. There is yet to elucidate a standard surveillance system for HCC recurrence risk owing to complexity of this malignancy. Generation of multi-omics data from patients facilitate the identification of robust signatures for various diseases. Thus, the current study is an attempt to develop the prognostic models employing multi-omics data to significantly (p-value <0.05) stratify the recurrence high-risk (median Recurrence Free Survival time (RFS) =<12 months) and low-risk groups (median RFS >12 months). First, we identified key 90RNA, 50miRNA and 50 methylation features and developed prognostic models; attained reasonable performance (C-Index >0.70, HR >2.5), on training and validation datasets. Subsequently, we developed a prognostic (PI) model by integrating the four multi-omics features (*SUZ12*, hsa-mir-3936, cg18465072, and cg22852503), that are biologically inter-linked with each other. This model achieved reasonable performance on training and validation dataset, i.e. C-Index 0.72, HR of 2.37 (1.61 - 3.50), p-value of 6.72E-06, Brier score 0.19 on training dataset, and C-Index 0.72 (95% CI: 0.63 - 0.80), HR of 2.37 (95% CI: 1.61 - 3.50), p-value of 0.015, Brier score 0.19 on validation dataset. Eventually, Drugbank data was investigated to elucidate therapeutic potential of these signatures. We have identified nine potential drugs against three genes (*CA9, IL1A, KCNJ15*) that are positively correlated with the tumor recurrence. We anticipate these results from our study will help researchers and clinicians to improve the HCC recurrence surveillance, eventually outcome of patients.

## 1. Introduction

HCC is among the most lethal malignant tumors that alone accounts for nearly 8.2% of the cancer-related deaths. Several factors like the timing of tumor detection, the severity of the disease condition, treatment modalities and tumor recurrence, etc. affect the survival rate of the patients [1–4]. Thus, optimal treatment and management of cancer could enhance the survival of patients. Over the last 2-3 decades, treatment strategies for HCC have been evolved, and their implementation depends upon a variety of factors including the size of the tumor, the pathogenesis of malignancy, feasibility of treatment, functioning, and performance of the liver [5–10]. For instance, liver resection surgery or liver transplantation (LT) or local ablation is recommended for the smaller tumor. LT depends upon the availability of the donor, liver resection surgery, and local ablation are the most commonly used treatments in the scenario [9–11]. Further, local ablation therapeutic options include microwave ablation (MWA),radiofrequency ablation (RFA), and cryoablation (Cryo-A) as the best treatment options for patients with BCLC stage 0-A, some BCLC stage B with TACE, especially where resection and LT are not suitable [10,12,13]. For large tumors, treatment options become limited, and the choice of treatment is still debatable. Literature indicated that the clinical outcome of patients with surgical resection is better than local ablation treatment, specifically for the patients having a well-preserved hepatic function[11]. However, postoperative tumor recurrence is one of the major concerns associated with the management and treatment of HCC patients. Tumor recurrence can be classified as early recurrence (ER) and late recurrence (LR) based on the period of recurrence post-surgery [14]. For instance, in early recurrence, intrahepatic new tumor lesions occur within 2 years of post-operation of HCC, while new intrahepatic lesions that occur after two years of surgery are categorized as late recurrence [14,15]. Patients with late recurrence have improved survival rates than that of early recurrence [14]. Overall, tumor recurrence drastically affects the prognosis of HCC patients. Therefore, the identification of patients who are at high risk of postoperative tumor recurrence is a crucial step to facilitate clinicians to provide appropriate curative therapy. Eventually, timely detection of tumor recurrence in patients and stratification of risk groups of the patients can enhance the prognosis of patients.

To date, there are no standard surveillance systems available that are employed for the stratification of high and low-risk groups of tumor recurrence in HCC patients. Although, the AJCC, TNM, and the BCLC staging methods based on the pathological information are used for the treatment regimens for HCC patients. But, their performance in recurrence risk stratification has not been evaluated appropriately [16]. Besides, in the recent past, some of the prognostic models designated for the prediction of tumor recurrence after surgical resection include Singapore Liver Cancer Recurrence (SLICER) score [17], the Nomogram Korean model [18], Surgery-Specific Cancer of the Liver Italian Program (SS-CLIP) [19]. The lack of validation on any independent dataset is a primary limitation associated with them [16–19]. Further, Yuan C et al. developed radiomics model based on portal venous phase with clinicopathological factors attained C-index 0.75 on the validation dataset [13]. Although this model performed well, the validation cohort contains 55 samples only [13]. Thus, to establish any inference from this small dataset is still debatable. Moreover, the number of studies revealed the association between different clinicopathological factors like AFP level, tumor size, HBsAg and vascular invasion, etc. with the tumor recurrence in HCC; thus, these were also established as prognostic indicators for prediction of HCC recurrence [13,16,20–23]. However, the molecular and genetic aspect of tumor recurrence remains invincible from them.

The advancement of omics and biomedical techniques generated an ample amount of genomic data from the patients and deposited in the public repositories like the GEO [24], TCGA [25], etc. This indeed facilitates the oncological researchers in the identification of genetic signatures or biomarkers for a variety of malignancies [26–47]. Thus, omics data can provide a better perspective regarding the pathways that derive tumor recurrence. In this regard, recently, Kong J et al. explored the transcriptomic data of patients for the elucidation of signature for recurrence prediction. They designated three genes including *PZP, SPP2*, and *PRC1* based signature for the recurrence risk prediction in patients and attained C index of 0.66 on training dataset and C index of 0.57 and 0.59 on two validation datasets [48]. As HCC is complex disease, a single omics layer might not be sufficient to understand the malignancy transformation [49–51]. Thus, multi-omics signatures need to be explored for risk prediction of HCC recurrence, which might provide a better option in this regard. Moreover, the integration of multi-omics layers can also enhance the performance of the prediction models, as reported in our previous study [52].

Thus, current study is an attempt to seek the potential of multi-omics data types, i.e., methylation (epigenomics), and miRNA in addition to transcriptomics data and their integration, for risk stratification in the context of tumor recurrence in HCC patients. Firstly, the critical omics features from each individual omics data type elucidated to predict tumor recurrence risk groups by employing the univariate survival analysis and various feature selection techniques. Here, we identified key feature sets (90 RNA, 50 miRNA and 50 methylation CpG sites) from three omics data types. Subsequently, prediction models developed to predict the labels of samples implementing machine learning algorithms. Eventually, the performance of models was evaluated on validation datasets. Further, these multi-omics features were also integrated and prognostic models were developed and validated.

## 2. Material and methods

### 2.1 Collection of Datasets

First, we obtained RNA-seq, miRNA-seq, and Methylation data for TCGA-LIHC patients from the GDC data portal [53] with the GDC-client tool. Next, Biospecimen, clinical, and metadata files are downloaded to extract clinical and demographic information of the patients by matching the TCGA bar code. Initially, we have 374 patient samples for RNA-seq and methylation data, while 369 samples for miRNA-seq. Notably, there were a total 60,483 RNA transcripts in RNA-seq data for each sample, and each transcript is quantified in terms of FPKM (fragments per kilobase of transcript per million mapped reads) value. miRNA-seq data contains FPKM value for 1,881 miRNAs. Further, there are 485,577 methylation CpG sites (Probe IDs) for each patient, and there is a methylation beta value corresponds to each CpG site. Besides the TCGA data, for external validation, we have also downloaded raw files for GSE14520 [54,55] and GSE6857 [56,57] datasets from the GEO using the GEOquery package of the Bioconductor [58,59]. Clinical and demographic data of patients was obtained from the series matrix files and respective publications [54–56]. GSE14520 dataset is the RNA-seq data that contains 22,268 probes for each of 221 HCC tumor samples. While GSE6857 is miRNA-seq data that contains 11,520 probes for each of the 193 HCC tumor samples. The distribution of samples in each dataset used in the study is represented in Figure S1 (Supplementary File 2).

### 2.2 Pre-processing of Datasets

Here, we have two datasets each for RNA-seq and miRNA-seq, and one dataset for methylation. First, we compared common samples among RNA-seq, miRNA-seq,and methylation profiles in the TCGA. We found there are 367 common samples among RNA-seq, miRNA-seq, and methylation data. For RNA-seq data, Entrez transcript IDs were mapped to the gene symbols based on the GENCODE v22 annotation file. Eventually, we obtained 19,831 protein-coding genes. In the methylation data, we have obtained a total of 485,577 methylation CpG sites. We have observed that there are missing values for methylation CpG sites among several samples. Hence, we excluded nearly 23-25% CpG sites, where beta value is missing among any of the samples employing in-house bash script. Eventually, we were left with 369,221 CpG sites for analysis in the study. For the external validation dataset, GSE14520 has raw CEL files of Affymetrix profiling techniques; these raw files are preprocessed using the Oligo package of Bioconductor [60,61], where background correction was followed by conversion into RMA values. Eventually, we have processed data with RMA values corresponding to 22,268 probes. Probe IDs were mapped to gene symbols using the respective platform files. Average of multiple probes correspond to single gene/miRNA computed to quantify the expression of single entity (Gene or miRNA) for each dataset using an in-house R script. Eventually, the GSE14520 contains 13,768 protein-coding genes for each of 221 samples, and the GSE6857 contains 479 unique miRNAs for each of 193 samples. On comparing, GSE14520 and GSE6857 datasets, we found 190 common samples among them. Eventually, we included only these 190 common samples for analysis. Clinical and demographic characteristics of patients are given in Table S1 (Supplementary File 1).

### 2.3 Partition of datasets

We have divided each of these datasets into 80:20 ratios, where 80% dataset is considered as a training dataset, and the remaining 20% of the dataset is used as an independent validation dataset. Thus, each of the training datasets contains 293 samples, and the validation dataset contains 74 samples. Further, we have observed that information related to recurrence-free survival is missing for some of the samples in training and validation datasets. Therefore, eventually, we are left with the training dataset containing 255 samples, and the validation dataset containing 57 samples for recurrence related analysis. Besides, GSE14520 and GSE6857 are used as external validation dataset.

### 2.4 Normalization of datasets

TCGA-RNA and miRNA-seq have FPKM values corresponding to each transcript or miRNA. We observed there is a wide range of variation in FPKM values; thus, first, we transformed FPKM values to log2 values after the addition of 1 as a constant to each value applying Equation 1 using in-house Rscript.

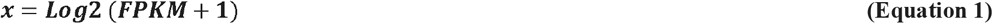

Where x is log 2 transformed value of the given gene, subsequently, data is quantile normalized using a training dataset as the target matrix employing the PreprocessCore package [62,63]. Besides, external validation datasets were also quantile normalized utilizing the training dataset as the target matrix. To develop the prediction models based on the integration of multi-omics data, we performed the Min-Max normalization (value between 0 to 1) method from the *caret* package in R [64].

### 2.5 Elucidation of Recurrence-related Features

Here, different steps were followed to extract the recurrence-related features, as represented in Figure 1. A detailed description of each step is in the following sections.

**Figure 1:**
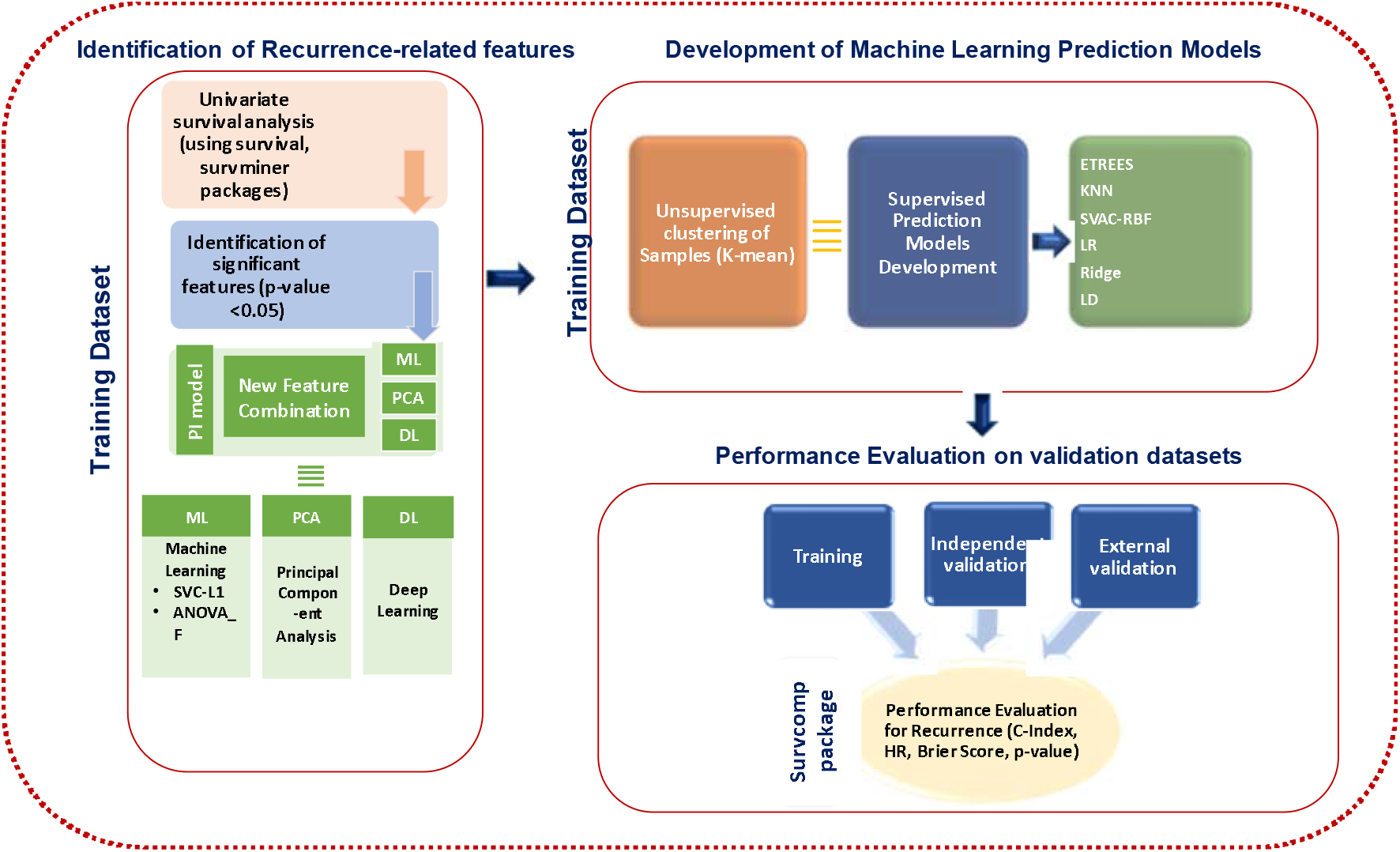
Overall workflow of this study representing various steps: feature selection, prediction model development on the training dataset, and their evaluation on independent and external validation datasets.

#### 2.5.1 Univariate survival analysis

Intending to scrutinize the features (RNA, miRNA, and methylation CpG sites) that are associated with the tumor recurrence, we performed univariate survival analysis for each feature using “survival” [65] and “survminer” [66] packages in R. Subsequently, we have selected only those features that stratified high-risk and low-risk groups with the p-value less than 0.05 implementing the log-rank test.

#### 2.5.2 Feature selection and extraction techniques

Based on the univariate survival analysis, we have obtained a large number of features. To further reduce the feature dimension from these significant features, we applied SVC-L1, ANOVA_F (or f_classif), principal component analysis (PCA), and autoencoder-based deep learning techniques. In our study, we used the SVC-L1 feature selection method, which was previously used for classification purposes in different malignancies conditions [45,67]. SVC-L1, ANOVA_F implemented using the Scikit python library [68]. In the SVC-L1, first, SVC selects the non-zero coefficients, and subsequently, the L1 penalty is applied to filter only relevant features from the data. Here, we also applied theANOVA_F feature selection method using the sklearn.feature_selection.f_classif function from the “scikit” package. It filters the vital features by computing the F-statistics. Among the feature extraction methods, we employed principal component analysis (PCA), and autoencoder-based deep learning (DL) techniques. While implementing PCA in R, we filtered only those components as features, which explained at least 1% of the variance in the data. The autoencoder-based deep learning method is implemented with three hidden layers (500, 100, and 500 nodes, respectively) using the Python Keras library [69] as adopted by many past studies [70,71].New 50 features were generated from the omics data by employing the bottleneck layer of the autoencoder.

#### 2.5.3 Prediction models

Here, we have used both unsupervised and supervised techniques to predict the labels of the samples based on the selected features. K-mean clustering is employed as an unsupervised machine learning technique that was implemented using python. Different supervised algorithms include KNN, SVC-RBF, ExtraTrees, LR (Logistic Regression), Ridge, LDA, etc. implemented using the Scikit-learn package[68]. Prediction models were developed by employing the ten-fold cross-validation technique on the training dataset.

#### 2.5.4 Performance Assessment

To estimate the performance of the in silico models developed to predict the labels of the samples, we have used several parameters like concordance index, Log-rank P-value, the Hazard ratio (HR), and Brier score that were previously used in different studies [70,71].

## 1. Results

### 3.1 Risk-stratification with Demographic and Clinical Factors

In the past, many studies revealed the role of demographic and clinical factors in the prognosis of liver cancer [1–4]. Thus, we explored three common factors, i.e., age, gender, and tumor stage, that are present among the training, independent validation, and external validation datasets for stratification of risk groups. Among them, the tumor stage was found to be a significant prognostic factor, as given in Table S2 (Supplementary File 1). Although, tumor stage is unable to stratify recurrence risk groups in independent validation dataset significantly, still, our analysis showed that tumor stage is an only parameter that significantly (with a p-value less than 0.0001) stratified recurrence risk groups of training and external validation dataset with C-Index 0.72 and 0.68, and attained HR 1.71 and 1.76, as represented in Figure S2 (Supplementary File 2).

### 3.2 Risk-stratification with Multi-omics Data

From the above analysis, we observed the tumor stage is an only significant prognostic factor for tumor recurrence prediction. Accurate stage prediction is another trivial task. This prompts us to explore omics features, i.e., RNA, miRNA, and methylation, to scrutinize vital features that can be explored for tumor recurrence risk stratification. There are a large number of features for each type of omics layer in the training dataset (RNA, miRNA, and methylation). For instance, there are 19,831 protein-coding genes, 1,881 miRNA, and 369,221 methylation CpG sites. Thus, to reduce the features, first, we applied univariate survival analysis concerning recurrence-free survival (RFS). Here, we obtained 1,560 protein-coding RNA, 66 miRNAs, and 10,394 methylation CpG sites that can significantly (p-value <0.05) stratified risk groups. Further, we also applied various feature selection techniques to filter variables and, consequently, developed ML prediction models based on the selected sets of features to predict the labels of samples. Prediction models and their performance are described in detail in the following sections.

#### 3.2.1 RNA-based prognostic models

Based on the univariate survival analysis, we obtained 1,560 protein-coding RNAs that are significantly associated with the RFS. We have clustered samples of training and validation datasets based on these features employing the k-means clustering approach. We have obtained C-Index 0.71 and 0.76 on the training and validation dataset, respectively. Out of 1,560 RNAs, 1,014 RNAs were found to be common among training and external validation dataset. Thus, we developed prediction models employing these 1,014 RNA features. We observed that in most cases, performance remains almost the same on training data; however, it decreased on the validation dataset. For instance, the LR model obtained a C-Index of 0.72, 0.62, and 0.58 on training, validation, and external validation dataset with significant p-value and with HR 3.11, 2.01, and 1.38, as shown in Table S3 (Supplementary File 1). Other models performed poor on the external validation dataset. The predictions by k-mean clustering achieved much higher performance, i.e., 0.78 and 0.86 on both training and validation dataset, respectively; but, drastically decreased on the external validation dataset. However, we did not get satisfying performance on an external validation dataset with 1,014 features. Hence, further, we employed various feature extraction and feature selection techniques including PCA, SVC-L1, autoencoder deep learning, and ANOVA_F to filter different feature sets like 28 by PCA, 90 by SVC-L1, 50 DL, and 100 ANOVA_F features. Subsequently, labels of samples were predicted employing different ML techniques. Among them, K-mean clustering prediction based on 28 components extracted by PCA is a top performer that attained C-Index of 0.77, 0.86 and 0.66, and HR of 2.49, 3.06 and 1.79 on the training, validation, external validation datasets, respectively with significant p-values, as shown in Table S4 (Supplementary File 1). Besides, other ML models based on these 28 features also achieved good performance on training, validation, and external validation dataset, as shown in Table S4 (Supplementary File 1). Kaplan-Meier plot depicts the stratification of risk groups, as shown in Figure S3 (Supplementary File 2). Further, 90 RNA features were selected by the SVC-L1 method. The univariate survival analysis indicates their significant prognostic potential on training data, as shown in Table S5 (Supplementary File 1). Out of these 90 features, 44 features are favourable factors for recurrence-free survival, as indicated by HR <1.0; whereas, 46 features were observed as unfavorable with HR >1.0. In addition, ML prediction models developed based on the combination of these 90 features; predictions by K-means clustering, ETREES, Ridge classifiers achieved significant performance in the stratification of recurrence risk groups, as tabulated in the Table 1. Kaplan-Meier plots (Figure 2) depict the stratification of recurrence risk groups by K-means clustering. This model significantly (p-value <0.01) stratified recurrence risk groups and achieved C-Index of 0.76, 0.85, and 0.66 with HR of 2.91, 3.65, and 1.79 on training, independent validation and external validation datasets, respectively. Further, it has been observed that median RFS is 8.68 months (264 days), 11.41 months (347 days), and 23.6 months(717.83 days) for high-risk groups and median RFS is 17.66 months (537 days), 26.65 months (810.5 days), and 40.1 months (1219.71 days) for low-risk groups of training, independent and external validation cohorts.

**Table 1:**
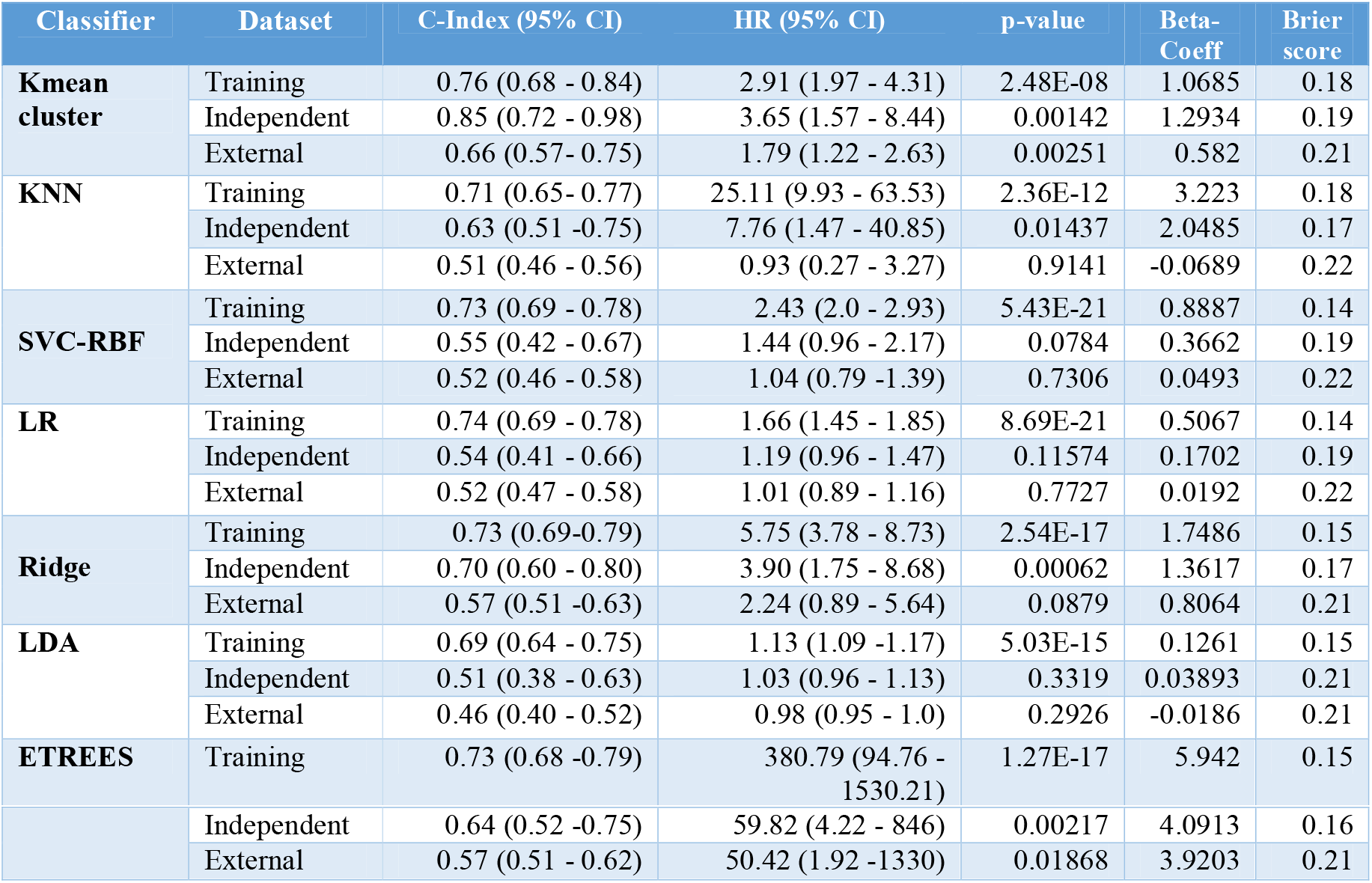
Performance of recurrence risk-prediction models based on 90 RNA features on training and validation datasets.

**Figure 2:**
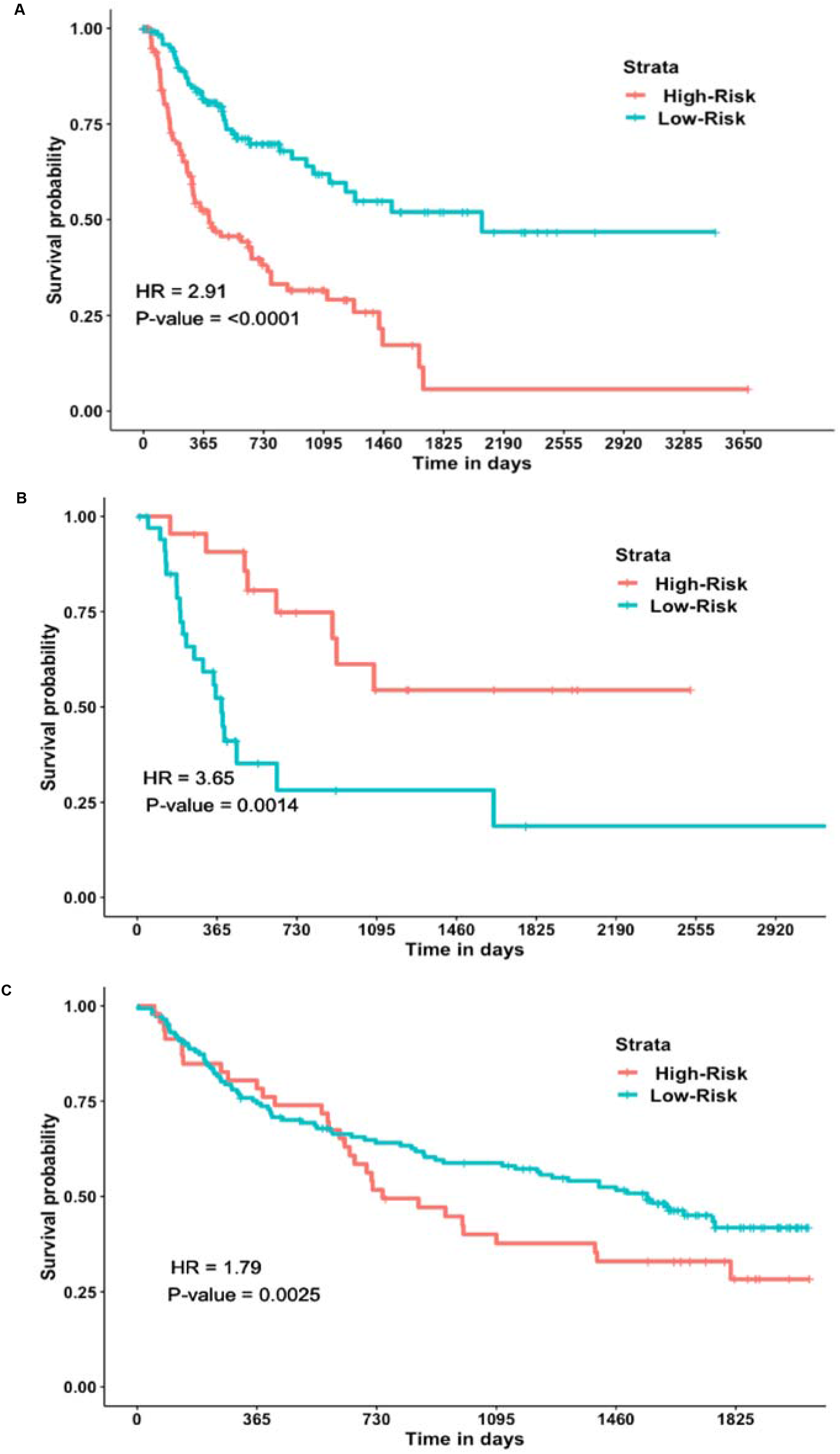
Kaplan-Meier plots illustrating the stratification of recurrence risk groups employing K-mean clustering predictions based on 90SVC-L1 RNA features: (A) training, (B) Independent, and (C) External validation dataset.

Further, to check whether the other feature sets enhance the performance. We have developed prediction models employing the 100 RNA features selected by ANOVA_F. Although the number of features increased in comparison to PCA and the SVC-L1, there is still a substantial decrease in the performance on independent and external validation datasets in most of the prediction models, as shown in Table S6(Supplementary File 1). In the recent past, few studies showed the improvement in the survival prediction models with the extraction of omics features with the autoencoder deep learning approach [70,71]. Thus, we also attempted an autoencoder deep learning (DL) approach for feature extraction. Here, we generated 50 features. Out of them, only 17 features were found to be significantly associated with the DFS by employing univariate survival analysis on these 50 features. The performance of the prediction models using these 17 features substantially decreased on the training as well as the validation dataset, as shown in Table S7(Supplementary File 1). In conclusion, based on the transcriptomics layer, we have obtained reasonable performance with 90 SVC-L1 feature sets and 28 PCA features set on training, independent validation, and external validation dataset. Out of them, the prediction based on k-mean clustering is the top performer.

#### 3.2.2 miRNA-based prognostic models

Next, miRNA expression data were explored for the identification of miRNA-signatures for recurrence risk stratification of HCC patients. Here, first, we obtained 66 miRNAs that were significantly stratified high-risk and low-risk recurrence groups of the training datasets based on univariate survival analysis employing recurrence-free survival. Subsequently, various prediction models were developed for the estimation of labels of samples. The external validation dataset contains only 10 miRNA out of 66 miRNA; thus, we validated most of our models only on an independent validation dataset. Prediction models based on these 66 miRNAs obtained reasonable performance on training and independent validation datasets. For instance, predictions based on k-mean clustering achieved the C-Index of 0.7 and 0.79 and HR 2.19, and 3.14 on the training and independent validation dataset with Brier score 0.2 and 0.19, respectively with significant p-value, as shown in Table S8 (Supplementary File 1). Ridge, SVC-RBF also performed quite well. Ridge achieved a C-Index of 0.69 and 0.71 and HR 3.37, and 11.17 on the training and independent validation dataset with Brier score 0.17 and 0.18.

Based on RNA transcripts, a prediction model that was developed employing 28 PCA components is the top performer. Thus, further, we extracted 31 features using PCA analysis that explains atleast 1% variation in the data. Subsequently, prediction models were developed by employing these features. Most of the prediction models developed using the 31 PCA features attained considerable performance in the estimation and stratification of recurrence risk groups. LR and Ridge models obtained C-Index of 0.69 and 0.72, HR nearly 4 and 12 with Brier score 0.18 and 0.17 on training and validation dataset, respectively, as represented in Table S9 (Supplementary File 1). Other models also achieved performance in a similar range.

Further, 50 miRNA features were selected by ANOVA_F. The univariate survival analysis revealed their significant potential for stratification of recurrence risk groups of training cohorts, as shown in Table S10 (Supplementary File 1). Among them, 16 were favorable and 34 unfavorable for recurrence-free survival with HR <1.0 and HR >1.0, respectively. Besides, we also attempted to develop prediction models based on these 50 miRNA features. Performance almost remains the same in most of the prediction models. There is a marginal improvement in the performance with the K-mean clustering prediction, i.e., achieved C-Index of 0.74, and 0.77 with HR of 2.61, and 2.94 on training, and independent validation datasets, respectively. Complete results are tabulated in Table 2. Kaplan-Meier plots (Figure 3) depict the stratification of recurrence risk groups by K-means clustering. Furthermore, we have observed the difference in median RFS time in both the risk groups, i.e., median RFS is 7.86 months (239 days), and 7.13 months (217days) for high-risk groups; while, median RFS is 15.68months (477 days), and 16.3 months (498days) for low-risk groups of training, and independent validation cohorts, respectively.

**Table 2:**
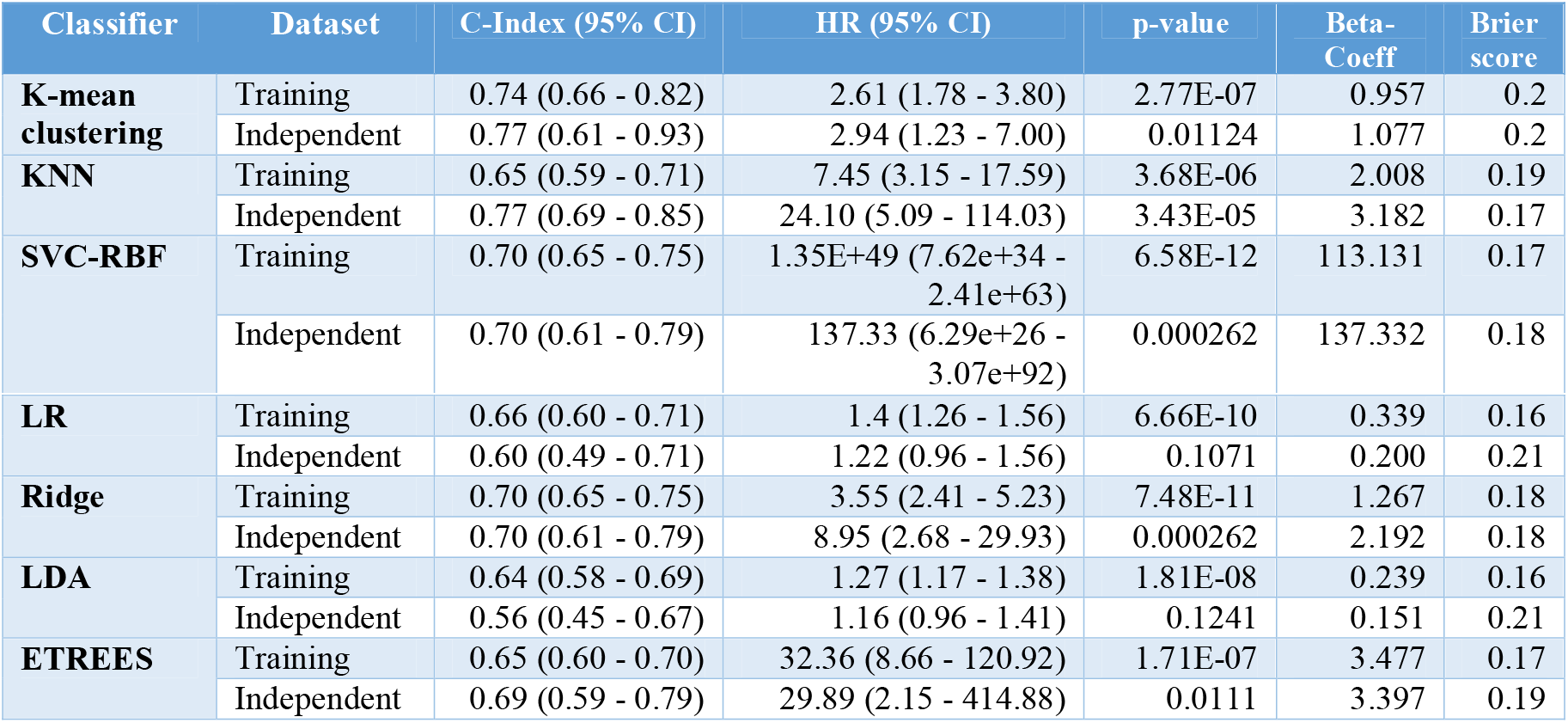
Performance of recurrence risk-prediction models based on 50 miRNA features on training and validation datasets.

| Classifier | Dataset | C-Index (95% CI) | HR (95% CI) | p-value | Beta-Coeff | Brier score |
| --- | --- | --- | --- | --- | --- | --- |
| **K-mean clustering** | Training | 0.74 (0.66 - 0.82) | 2.61 (1.78 - 3.80) | 2.77E-07 | 0.957 | 0.2 |
|  | Independent | 0.77 (0.61 - 0.93) | 2.94 (1.23 - 7.00) | 0.01124 | 1.077 | 0.2 |
| **KNN** | Training | 0.65 (0.59 - 0.71) | 7.45 (3.15 - 17.59) | 3.68E-06 | 2.008 | 0.19 |
|  | Independent | 0.77 (0.69 - 0.85) | 24.10 (5.09 - 114.03) | 3.43E-05 | 3.182 | 0.17 |
| **SVC-RBF** | Training | 0.70 (0.65 - 0.75) | 1.35E+49 (7.62e+34 - 2.41e+63) | 6.58E-12 | 113.131 | 0.17 |
|  | Independent | 0.70 (0.61 - 0.79) | 137.33 (6.29e+26 - 3.07e+92) | 0.000262 | 137.332 | 0.18 |
| **LR** | Training | 0.66 (0.60 - 0.71) | 1.4 (1.26 - 1.56) | 6.66E-10 | 0.339 | 0.16 |
|  | Independent | 0.60 (0.49 - 0.71) | 1.22 (0.96 - 1.56) | 0.1071 | 0.200 | 0.21 |
| **Ridge** | Training | 0.70 (0.65 - 0.75) | 3.55 (2.41 - 5.23) | 7.48E-11 | 1.267 | 0.18 |
|  | Independent | 0.70 (0.61 - 0.79) | 8.95 (2.68 - 29.93) | 0.000262 | 2.192 | 0.18 |
| **LDA** | Training | 0.64 (0.58 - 0.69) | 1.27 (1.17 - 1.38) | 1.81E-08 | 0.239 | 0.16 |
|  | Independent | 0.56 (0.45 - 0.67) | 1.16 (0.96 - 1.41) | 0.1241 | 0.151 | 0.21 |
| **ETREES** | Training | 0.65 (0.60 - 0.70) | 32.36 (8.66 - 120.92) | 1.71E-07 | 3.477 | 0.17 |
|  | Independent | 0.69 (0.59 - 0.79) | 29.89 (2.15 - 414.88) | 0.0111 | 3.397 | 0.19 |

**Table 3:**
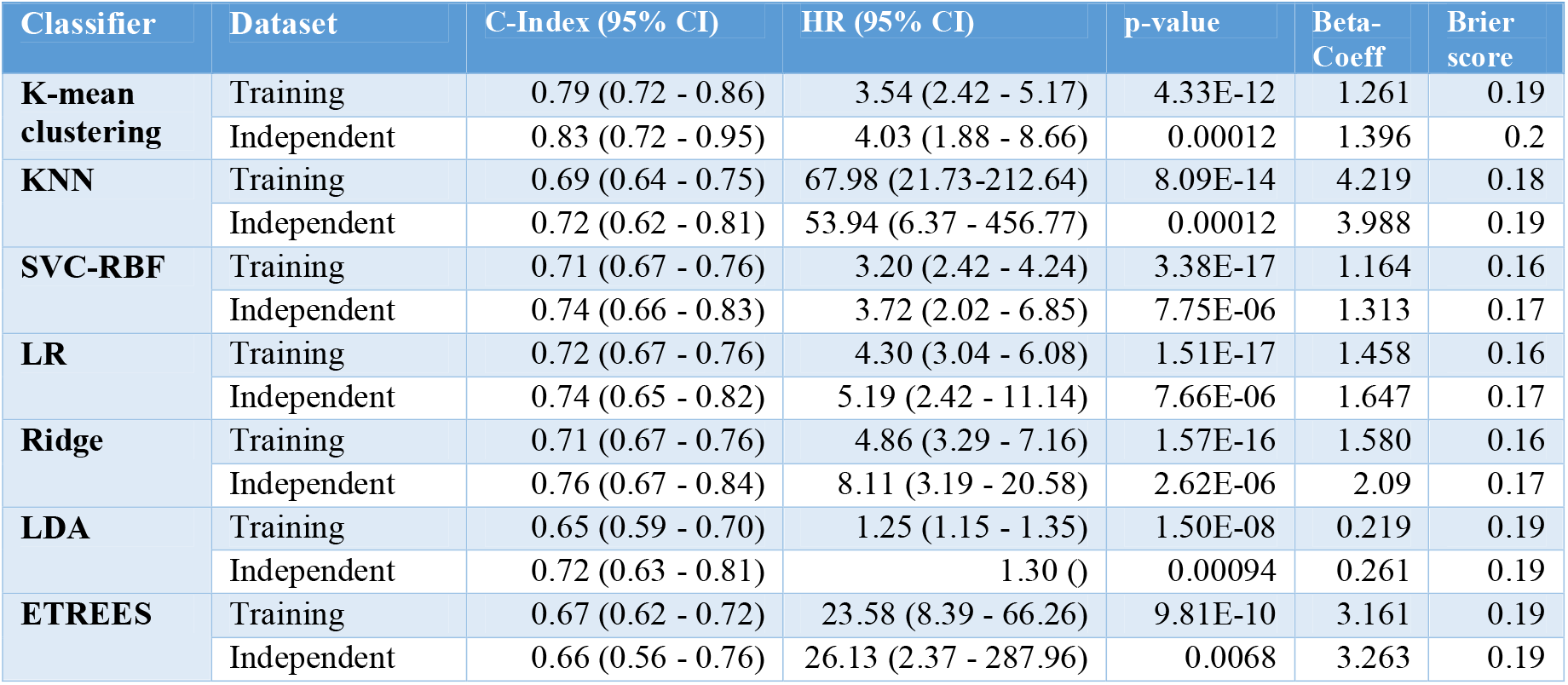
Performance of recurrence risk-prediction models based on 50 methylation CpG sites features on training and validation datasets.

| Classifier | Dataset | C-Index (95% CI) | HR (95% CI) | p-value | Beta-Coeff | Brier score |
| --- | --- | --- | --- | --- | --- | --- |
| **K-mean clustering** | Training | 0.79 (0.72 - 0.86) | 3.54 (2.42 - 5.17) | 4.33E-12 | 1.261 | 0.19 |
|  | Independent | 0.83 (0.72 - 0.95) | 4.03 (1.88 - 8.66) | 0.00012 | 1.396 | 0.2 |
| **KNN** | Training | 0.69 (0.64 - 0.75) | 67.98 (21.73-212.64) | 8.09E-14 | 4.219 | 0.18 |
|  | Independent | 0.72 (0.62 - 0.81) | 53.94 (6.37 - 456.77) | 0.00012 | 3.988 | 0.19 |
| **SVC-RBF** | Training | 0.71 (0.67 - 0.76) | 3.20 (2.42 - 4.24) | 3.38E-17 | 1.164 | 0.16 |
|  | Independent | 0.74 (0.66 - 0.83) | 3.72 (2.02 - 6.85) | 7.75E-06 | 1.313 | 0.17 |
| **LR** | Training | 0.72 (0.67 - 0.76) | 4.30 (3.04 - 6.08) | 1.51E-17 | 1.458 | 0.16 |
|  | Independent | 0.74 (0.65 - 0.82) | 5.19 (2.42 - 11.14) | 7.66E-06 | 1.647 | 0.17 |
| **Ridge** | Training | 0.71 (0.67 - 0.76) | 4.86 (3.29 - 7.16) | 1.57E-16 | 1.580 | 0.16 |
|  | Independent | 0.76 (0.67 - 0.84) | 8.11 (3.19 - 20.58) | 2.62E-06 | 2.09 | 0.17 |
| **LDA** | Training | 0.65 (0.59 - 0.70) | 1.25 (1.15 - 1.35) | 1.50E-08 | 0.219 | 0.19 |
|  | Independent | 0.72 (0.63 - 0.81) | 1.30 () | 0.00094 | 0.261 | 0.19 |
| **ETREES** | Training | 0.67 (0.62 - 0.72) | 23.58 (8.39 - 66.26) | 9.81E-10 | 3.161 | 0.19 |
|  | Independent | 0.66 (0.56 - 0.76) | 26.13 (2.37 - 287.96) | 0.0068 | 3.263 | 0.19 |

**Figure 3:**
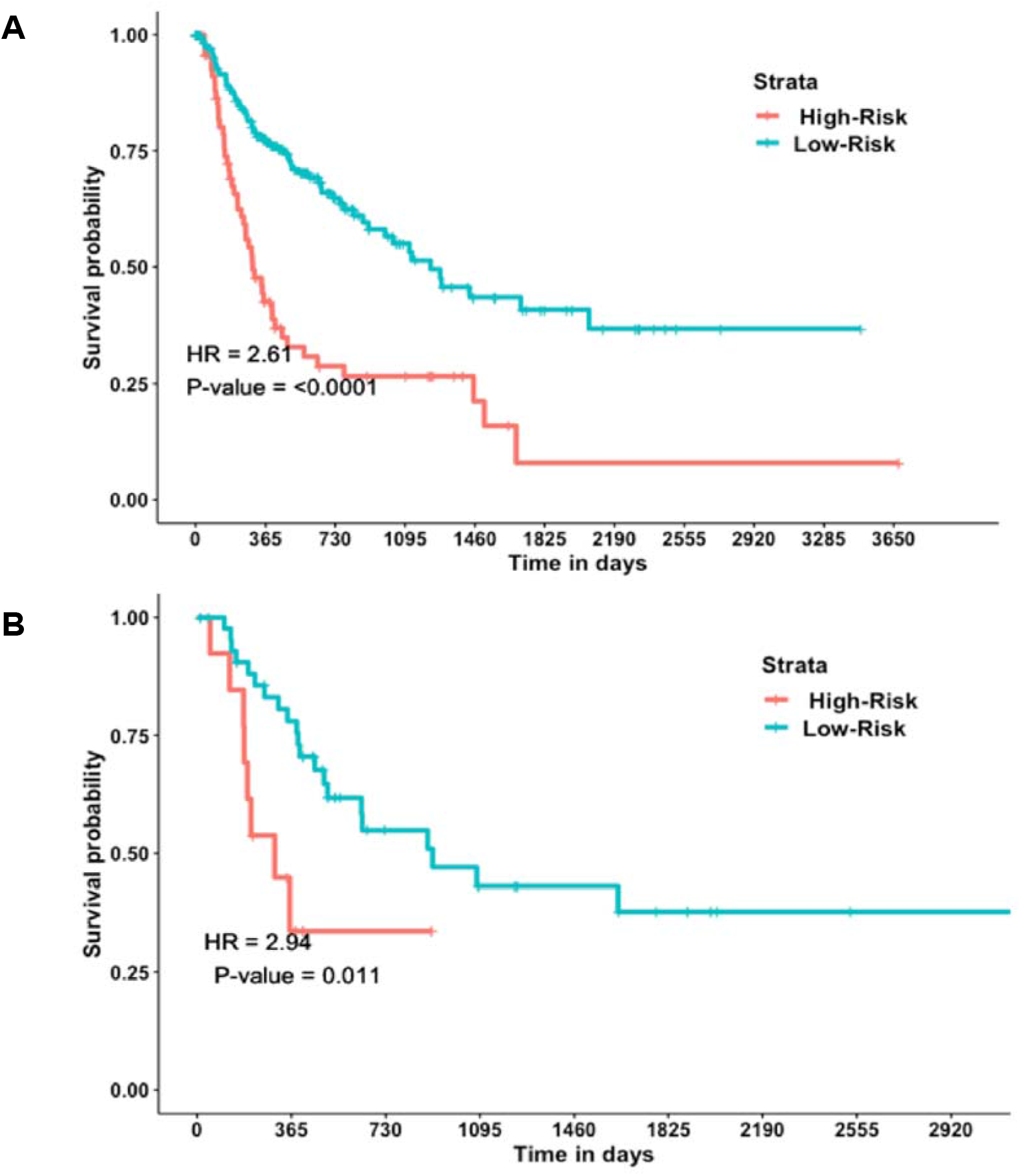
Kaplan-Meier plots depict the stratification of recurrence risk-groups with K-mean clustering predictions based on 50 miRNA features: (A) Training and (B) Independent validation Dataset.

As this feature set (50 miRNAs) performed quite well on training and independent validation dataset, next, we also attempted to validate this model on the external validation dataset. Unfortunately, external validation contains only 10 common miRNAs. Thus, we developed the model only on these 10 features. The performance drastically decreased on training as well as validation datasets (see Table S11, Supplementary File 1). Although, performance remains significant on the training dataset, but become insignificant on the validation datasets.

#### 3.2.3 Methylation-based prognostic models

To elucidate the importance of epigenomics patterns in the recurrence risk prediction, first, we performed univariate survival analysis on the CpG methylation data of training data. Thus, we obtained 10,394 CpG methylation sites that are significantly associated with the stratification of recurrence risk groups. Subsequently, labels of samples were predicted based on these features implementing various ML techniques. Although, most of these models obtained significant performance, quite lower than that of the prediction models developed based on the other two omics layers, i.e., RNA and miRNA, as shown in Table S12 (Supplementary File 1). Prediction models based on the 10,394 methylation CpG sites unable to achieve much higher performance. Thus, next, we select features from this set, employing different feature selection and extraction approaches similar to RNA and miRNA based models. First, 61 features were selected with SVC-L1. After that, prediction models developed based on these 61 features. Interestingly, despite the reduction in a substantial amount in the number of features, most of the prediction models attained higher performance than of models based on 10,394 methylation CpG sites. For instance, SVC-RBF, LR achieved C-Index more than 0.8 on the training dataset and 0.65 on the validation dataset. KNN and Ridge performed reasonably on the training dataset, i.e., C-Index nearly 0.75 on the training dataset and attained C-Index 0.69 and 0.67 on training and validation dataset, respectively. Complete results tabulated in Table S13 (Supplementary File 1).

Further, we selected 50 methylation CpG sites employing the ANOVA_F method. The univariate survival analysis of these features suggests 16 CpG sites are favorable factors for DFS, i.e., HR < 1.0; while, 34 features are unfavorable factors, i.e., HR >1.0 (see Table S14, Supplementary File 1). Further, ML prediction models developed based on the combination of these 50 features. The performance of most of the prediction models decreased on the training dataset, but it substantially improved on the validation dataset, as shown in Table 5. K-mean clustering prediction is a top performer that significantly (p-value <0.01) stratified recurrence risk groups and achieved C-Index of 0.79, and 0.83 with HR of 3.54 and 4.03 on training and independent validation datasets, respectively. Kaplan-Meier plots illustrate the stratification of tumor recurrence risk groups with the K-mean clustering based on these 50 methylation CpG sites, as shown in Figure 4. In addition, the difference in median RFS time between risk groups was observed, i.e., median RFS is 8.88 months (270 days), and 9.24 months (281days) for high-risk groups; while, median RFS is 15.68 months (477 days), and 17.59 months (535 days) for low-risk groups of training, and independent validation cohorts, respectively. Besides, similar to RNA and miRNA data, PCA (Table S15, Supplementary File 1) and DL also applied for the extraction of methylation-based features. But, the performance of models drastically decreased.

**Figure 4:**
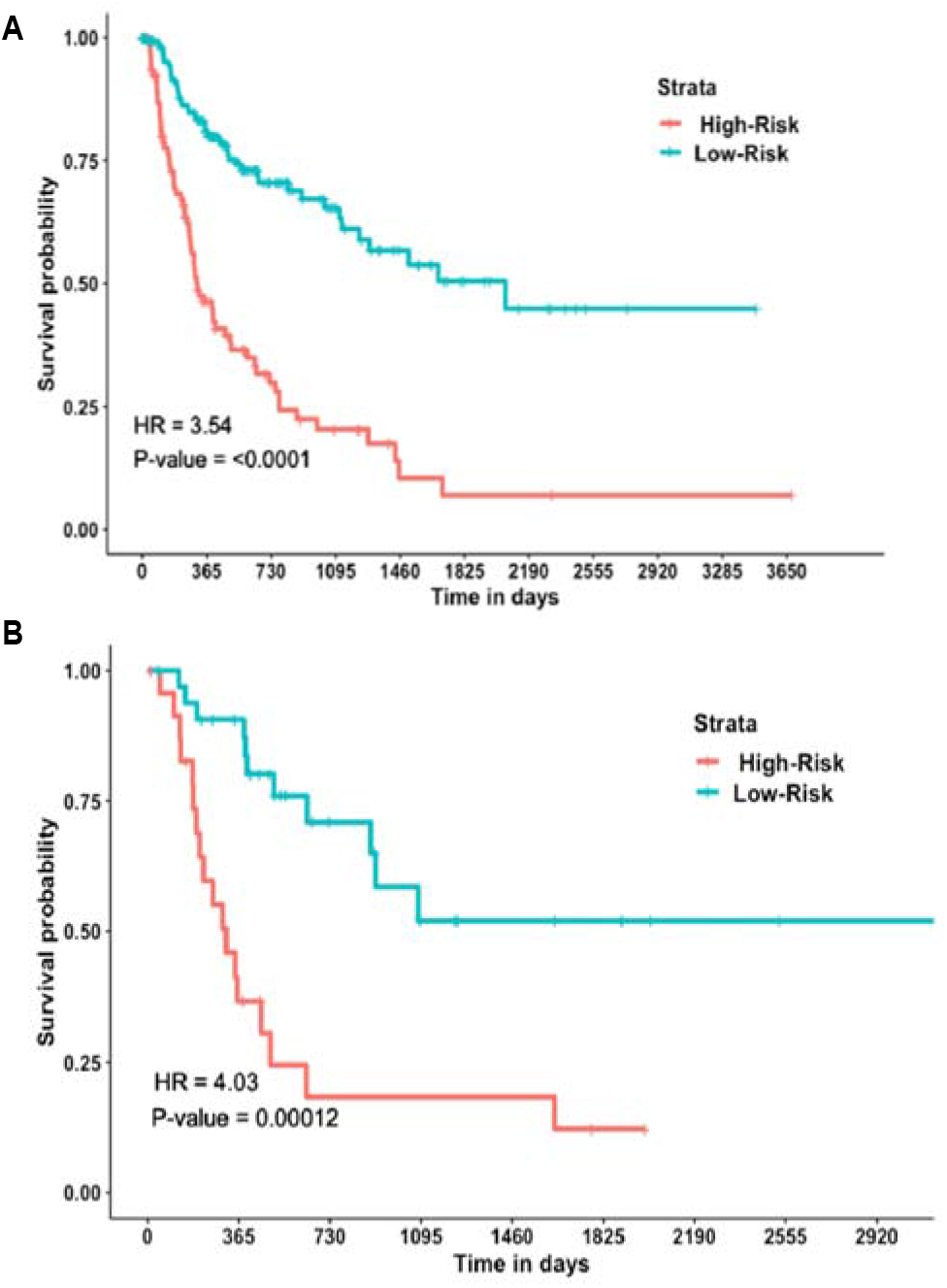
Kaplan-Meier plots illustrating the stratification of recurrence risk-groups with K-mean clustering predictions based on 50 methylation CpG sites: (A) Training and (B) Independent validation Dataset.

### 3.3 Prognostic models based on integration of multi-omics data

#### 3.3.1 Integrative models

Till now, we employed each omics layer individually to predict the recurrence risk groups. Next, we combined all the vital features from three omics layers, i.e., RNA, miRNA, and methylation CpG sites, to understand whether the integration of multi-omics layers improves the overall performance. For this, first, we combined key features that perform the best, i.e., 90 RNA, 50 miRNA, and 50 methylation CpG Sites. But, the performance of models is marginally decreased as compared to the models based on individual omics layers on both training as well as the validation datasets, as represented in Table 6. Predictions with K-mean clustering is the best performer, i.e., obtained C-Index 0.72 with HR of 2.22 on the training dataset and C-Index 0.80 with HR 2.92 on the validation dataset, respectively, as shown in Table S16 (Supplementary File 1). Moreover, we have observed the difference in median RFS time between high and low-risk groups, i.e., median RFS is 15.68 months (477 days), and 17.59months (535 days) for high-risk groups; while, median RFS is 16.44 months (500 days), and 16.6 months (505 days) for low-risk groups of training, and independent validation cohorts, respectively.

Further, we combined all the significant features obtained from univariate survival analysis that include 1560 RNA, 66 miRNA, and 10,394 methylation CpG sites. Subsequently, we extracted 10 PCA and 12 auto-encoder deep learning features and developed prediction models based on them. But performance is further decreased both on the training and validation dataset, as shown in Table S17 and Table S18 (Supplementary File 1).

#### 3.3.2 Prognostic index (PI) Model

Next, to find the unique set of multi-omics features that are biologically inter-linked with each other, we analyzed significant features (based on univariate COC-PH model) from all the three omics layers that include 1560 RNA, 66 miRNA, and 10,394 methylation CpG sites and their target. Towards this, we have identified target genes of significant miRNA candidates and using the miRWalk [72,73] and also scrutinized genes that are associated with significant methylation CpG sites. Subsequently, 1,014 RNAs, target genes of miRNA and methylation associated genes were compared. We observed hsa-mir-3936 targetone of the protein-coding genes, i.e., *SUZ12* gene, which is also present among 1560 RNAs. Besides, we found two methylation CpG sites, i.e., cg18465072, cg22852503 that are located on *SUZ12gene*, are among the 10,394 methylation significant methylation features. Thus, eventually, we developed prognostic Index (PI) model employing only these four features that inter-linked three omics layers (RNA, miRNA, methylation) of HCC patients;using following equation 2:

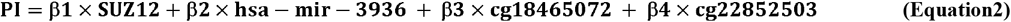

Where, β1=0.39, β2=0.48, β3=0.45 and β4=(−0.45) are the regression coefficient for *SUZ12*, hsa-mir-3936, cg18465072 and cg22852503, respectively, obtained using a univariate cox-PH model. Thus, the final PI equation becomes as follow:

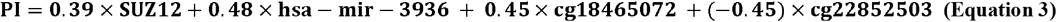

This PI model attained reasonable performance, i.e., significantly stratified risk groups of training dataset with C-Index 0.72, HR of 2.37 (1.61 - 3.50), a p-value of 6.72E-06, Brier score 0.19. This model almost performed in similar manner on independent validation dataset, i.e., C-Index 0.72 (95% CI: 0.63 - 0.80), HR of 2.37 (95% CI: 1.61 - 3.50), p-value of 0.015, Brier score 0.19. Kaplan-Meyer plot (Figure 5) representing the stratification of the group based on the PI model. The lack of hsa-mir-3936 in the external cohort and unavailability of external methylation data prohibits their validation on external cohorts. Further, we have observed the difference in median RFS time in both the risk groups, i.e., median RFS is 9.63 months (293 days), and 12.18 months (370.5days) for high-risk groups; while, median RFS is 15.73 months (478.5 days), and 16.6 months (505 days) for low-risk groups of training, and independent validation cohorts, respectively. Additionally, we have observed SUZ12 is significantly (Bonferroni p-value <0.01) upregulated in high-risk recurrence group in comparison to the low-risk recurrence group of training and independent validation dataset.

**Figure 5:**
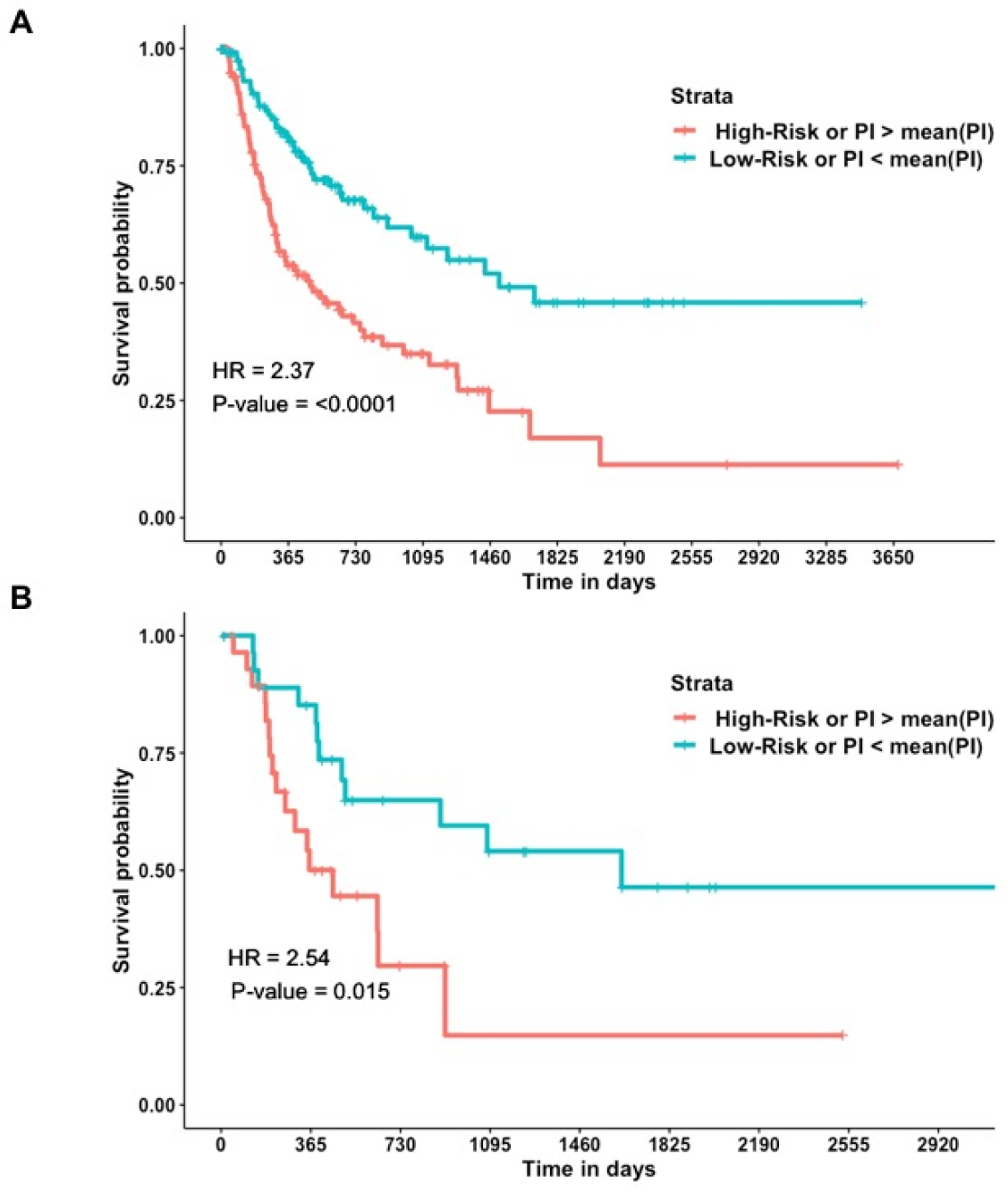
Kaplan-Meier plots representing the stratification of recurrence risk-groups with PI Model (hsa-mir-3936*0.48+ *SUZ12*\*0.39+ cg18465072*0.45+ cg22852503*(- 0.45)): (A) Training, and (B) Independent validation Dataset.

### 3.4 Biological Implications and Therapeutic Applications

*SUZ12* (SUZ12 Polycomb Repressive Complex 2 Subunit) is a Protein Coding gene. It is a component of the PRC2/EED-EZH2 complex, which methylates ‘Lys-9’ (H3K9me) and ‘Lys-27’ (H3K27me) of histone H3, leading to transcriptional repression of the affected target gene [74]. Further, we observed different studies reported SUZ12 for the treatment of hepatocellular carcinoma [75,76]. We have also tried to investigate the therapeutic applications of the identified prognostic markers. We first mapped the identified 66 prognostic miRNAs to the 1014 survival-related genes which resulted in 691 unique marker genes. Then, we searched for the existing drug targets against these genes using the DrugBank data (https://www.drugbank.ca/). We found the 511 active drug targeting 46 unique prognostic genes that were already approved for targeting these genetic markers in various diseases (refer Table S19, Supplementary File 1). When we investigated both active and inactive drugs against those 691 marker genes, it resulted in 2241 drugs against 146 unique genes (Table S20, Supplementary File 1). Further, we found a total of 178 prognostic CpG sites in 18 unique survival related genes (out of 1014). Drugs were then indentified for these survival-related genes with prognostic methylation sites. We found that only one survival related gene (*PTH2R*) with prognostic CpG site had pharmacologically active drug for it. In addition to that, three such unique genes namely *RNGTT, MAPK4* and *PTH2R* (CpG site related) had nearly eight active and inactive drugs designed against them (Table S21, (Supplementary File 1). Next, we focussed on drugs against 46 unfavourable genes (see Table S5, Supplementary File 1), i.e., whose expression positively correlated with tumor recurrence in HCC patients. Here, we found nine potential drugs against three genes, i.e., Rilonacept against *IL1A,* Yohimbine, and Butamben against *KCNJ15*, and Benzthiazide, Hydroflumethiazide, Zonisamide, Girentuximab, Ellagic acid, and Sodium carbonate against *CA9* gene. Complete results with drug ID and drug type are represented in Table S22 (refer Supplementary File 1). As these genes are positively correlated with tumor recurrence, thus their inhibition via these drugs might improve the overall prognosis of the HCC patients. We also tried to identify any available drugs for *SUZ12* which is functionally important being the intersecting gene with related miRNA and CpG sites. However, we could not find any active or inactive drug against this gene.

## 4. Discussion and Conclusion

The various therapeutic strategies like liver resection, transplantation, radiofrequency ablation, etc. improved the survival probability of HCC patients [77-80]. However, the late diagnosis and recurrence are among the key factors that are responsible for poor prognosis [1–4]. Hence, it is vital to elucidate the omics signature or biomarkers for tumor recurrence in HCC. The availability of omics data of liver cancer patients facilitates the research in discovering the robust signatures for liver cancer [52,81–83]; hence, multi-omics data can provide a holistic picture of this malignacy. Thus, in the present study, we have made an attempt to explore the multi-omics layers of HCC patients (obtained from public repositories, i.e., GDC data portal and GEO) for the elucidation of signatures that can significantly stratify the recurrence risk groups.

In the current work, first, we explored the demographic and clinical factors that include age, gender, and tumor stage for their prognostic potential in tumor recurrence. Our study revealed that the tumor stage is only a significant (p-value <0.0001) prognostic factor that stratified recurrence risk groups with C-Index 0.72 and 0.68 of training and external validation dataset. However, the performance on the independent dataset remains dismal. Besides, past literature indicates that accurate tumor stage prediction is itself a challenging task [45,52,67,84,85]. This propels us to analyze the multi-omics data of patients for the elucidation of recurrence related signatures. Towards this, first, we identified 1,560 RNAs, 66miRNAs, and 10,494 methylation CpG sites that are significantly associated with tumor recurrence in HCC, implementing the univariate survival analysis on the training dataset. Subsequently, different feature selection and extraction approaches such as SVC-L1, F-ANOVA, PCA, and auto-encoder deep learning (DL) were implemented to filter vital features for each omics layer individually. Consequently, various machine learning techniques include K-mean clustering, SVC-RBF, ETREES, LR, LDA, KNN, Ridge classifier, etc. employed to predict the labels or risk group of samples based on selected features. Thereafter, Cox-PH survival analysis was performed for the estimation of recurrence risk stratification.

Among the RNA data, we found 28 PCA features and 90 SVC-L1 RNA features that significantly stratified recurrence risk groups in training and validation datasets. Mainly, prediction models based on 28 PCA features are the top performers. For instance, predictions with k-mean clustering attained C-Index 0.77, 0.82, and 0.66 with a p-value less than 0.0001 on training, independent and external validation cohorts, respectively. Furthermore, we have identified 50 miRNA selected by ANOVA_F, prediction models based on them achieved quite good performance, i.e., C-Index 0.74 and 0.77 with a p-value less than 0.0001 on the training and independent validation dataset, respectively. In addition to RNA and miRNA signatures, our study found 50 methylation CpG sites that significantly classified recurrence risk groups. Like, K-mean clustering predictions based on these 50 methylation CpG sites achieved C-Index 0.79 and 0.83 with a p-value of less than 0.0001 on training and independent validation dataset, respectively. However, performance decreased on other ML prediction models. Overall, we observed that prediction with 28PCA from RNA features, 90 SVC-L1 RNA features, 50 miRNA, and 50 methylation CpG sites selected by ANOVA_F are the top performers and achieved performance almost in similar range on training and independent validation dataset. Next, we integrate features from the three omics layers and develop prediction models to understand whether the integration of multi-omics data enhances performance. We observed prediction models based on the integration of most vital features together, i.e., 90 RNA, 50 miRNA, and 50 methylation CpG sites, unable to enhance the performance further, as compared to the models based on individual omics data. Besides, we also attempted to develop prediction models by extracting the features using PCA and DL based on the integration of omics data, but the performance does not improve.

At last, we attempted to develop a prognostic model employing four features (*SUZ12*, hsa-mir-3936, cg18465072, and cg22852503) that are common and inter-linked between these multi-omics layers of HCC. We found *SUZ12* as a target of hsa-mir-3936 and cg18465072, and cg22852503 are two methylation CpG sites that are associated with *SUZ1*. Thus, we developed a prognostic Index (PI) model employing only these four features. We observed, this PI model significantly stratified high-risk (>12 months or 1 year) and low-risk (<= 12 months or 1 year) recurrence groups of HCC patients; achieved C-Index 0.72 on training and independent validation datasets.

*SUZ12* encodes a protein subunit of transcription Polycomb Repressive Complex (PRC2) that represses target proteins. We observed the significant upregulation of *SUZ12* in high-risk recurrence group. Previously, the activation of SUZ12/PRC2 complex was reported in liver carcinogenesis and HBV infection [86]. Besides, recently, overexpression of key components of PRC2, i.e., EZH2/SUZ12/EED, was found to be associated with poor prognosis of cholangiocarcinoma [87]. In addition, the upregulation of *SUZ12* was reported to be a prognostic factor for poor prognosis of various malignancies [88,89]. But, in contrast, recently, one of the studies has shown that the lower expression of *SUZ12* results in poor prognosis in terms of the 5-year survival of HCC patients [76]. Xue et al, further demonstrated the significant down-regulation of *SUZ12* in HBV-related HCC tissue samples in comparison to normal samples; and its overexpression impaired HCC cell’s migration and invasion [76]. Thus, more studies on HCC cohorts can confirm the exact role of *SUZ12* in the prognosis of HCC patients.

Although, the RNA based prediction model in the past that utilized three genes, i.e., *PZP, PRC1,* and *SPP2*, achieved C-Index0.66 on the training cohort and C-index of 0.57 and 0.59 on two independent validation datasets [48]. In our study, the performance of models has reasonably improved on the training and the validation data. Most of the previous studiesfocussed on transcriptomic databut, we explored three omics layers, i.e., RNA, miRNA, and methylation, to elucidate the vital prognostic candidates that include 90 RNA, 50 miRNA, and 50 methylation CpG sites. These significantly segregated the recurrence high-risk (median RFS time <= 12 months) and low-risk (median RFS time > 12 months)groups of training and validation datasets. To the best of authors’ knowledge, this is the first attempt, where the integrative multi-omics approach was explored for recurrence risk stratification of HCC patients. PI model based on four integrative multi-omics features (*SUZ12*, hsa-mir-3936, cg18465072, and cg22852503) significantly stratified risk groups of training and validation cohorts with C-Index 0.72.These multi-omics candidates can be further exploited to understand the underlying complex mechanisms of tumor recurrence from a holistic perspective. Additionally, we have also made an attempt to identify the drugs designed against the prognostic genes linked to survival related miRNAs and CpG sites, respectively. We found 551 drugs for 41 prognostic genes linked to miRNA survival-related sites, whereas 8 drugs against 3 unique marker genes with survival-related methylation sites. Overall, 559 drugs were identified targeting 49 different survival-associated genes. Our study also found nine potential drugs (Rilonacept, Yohimbine, Butamben, Benzthiazide, Hydroflumethiazide, Zonisamide, Girentuximab, Ellagic acid, and Sodium carbonate against three unfavourable genes (*IL1A, KCNJ15, CA9*). These drugs can be employed to devise the risk-group specific therapeutic strategies as explained above in detail. The drugs identified for marker genes above can be used to devise the expression based therapeutics to patients in the respective risk groups, specifically. As risk prediction was done on the basis of median cut-off values of the gene expressions, these known drug-target information can be exploited to design a more reliable and patients-risk group specific therapy which can eventually inhibit/activate the expression levels for the desired outcome.

Eventually, we anticipate this study would be beneficial for the researchers working in the area of liver malignancy to understand the recurrence risk groups stratification of HCC patients from a broader perspective, ultimately facilitates to improve the outcome of patients.

## Author Contributions

H.K. collected the data and created the datasets, H.K. and A.L. developed algorithms, H.K. implemented algorithms. H.K., A.L. and G.P.S.R. analyzed the results. H.K., A.L. wrote the manuscript. G.P.S.R. conceived and coordinated the project, helped in the interpretation and analysis of data, refined the drafted manuscript and gave complete supervision to the project. All of the authors have read and approved the final manuscript.

## Funding

This research was funded by J. C. Bose National Fellowship (with Grant No. SRP076), Department of Science and Technology (DST), INDIA.

## Acknowledgments

All the authors acknowledge funding agencies J. C. Bose National Fellowship (DST). H.K. and A.L. are thankful to Council of Scientific and Industrial Research (CSIR) and UGC, India for providing fellowships, respectively.

## Conflicts of Interest

The authors declare no financial and non-financial conflict of interest.

## Supplementary Data

Supplementary File 1: Supplementary Tables.

Supplementary File 2: Supplementary Figures.

## Notes

### Competing Interest Statement

The authors have declared no competing interest.

### Summary of Updates

Authors order updated.

## References

[1] S.K. Park, Y.K. Jung, D.H. Chung, K.K. Kim, Y.H. Park, J.N. Lee, O.S. Kwon, Y.S. Kim, D.J. Choi, J.H. Kim, Factors influencing hepatocellular carcinoma prognosis after hepatectomy: A single-center experience, Korean J. Intern. Med. 28 (2013) 428–438. https://doi.org/10.3904/kjim.2013.28.4.428.

[2] M.Y. Cai, F.W. Wang, C.P. Li, L.X. Yan, J.W. Chen, R.Z. Luo, J.P. Yun, Y.X. Zeng, D. Xie, Prognostic factors affecting postoperative survival of patients with solitary small hepatocellular carcinoma, Chin. J. Cancer. 35 (2016) 80. https://doi.org/10.1186/s40880-016-0143-x.

[3] M. Abdel-Wahab, T.S. El-Husseiny, E. El Hanafy, M. El Shobary, E. Hamdy, Prognostic factors affecting survival and recurrence after hepatic resection for hepatocellular carcinoma in cirrhotic liver, Langenbeck’s Arch. Surg. 395 (2010) 625–632. https://doi.org/10.1007/s00423-010-0643-0.

[4] A. Chiappa, A.P. Zbar, R.A. Audisio, B.E. Leone, F. Biella, C. Staudacher, Factors affecting survival and long-term outcome in the cirrhotic patient undergoing hepatic resection for hepatocellular carcinoma, Eur. J. Surg. Oncol. 26 (2000) 387–392. https://doi.org/10.1053/ejso.1999.0904.

[5] M.H. Attwa, S.A. El-Etreby, Guide for diagnosis and treatment of hepatocellular carcinoma, World J. Hepatol. 7 (2015) 1632–1651. https://doi.org/10.4254/wjh.v7.i12.1632.

[6] J.M. Llovet, J. Fuster, J. Bruix, The Barcelona approach: Diagnosis, staging, and treatment of hepatocellular carcinoma, Liver Transplant. 10 (2004) S115–S120. https://doi.org/10.1002/lt.20034.

[7] S.K. Olsen, R.S. Brown, A.B. Siegel, Hepatocellular carcinoma: Review of current treatment with a focus on targeted molecular therapies, Therap. Adv. Gastroenterol. 3 (2010) 55–66. https://doi.org/10.1177/1756283X09346669.

[8] A.V.C. França, J.E. Junior, B.L.G. Lima, A.L.C. Martinelli, F.J. Carrilho, Diagnosis, staging and treatment of hepatocellular carcinoma, Brazilian J. Med. Biol. Res. 37 (2004) 1689–1705. https://doi.org/10.1590/S0100-879X2004001100015.

[9] Z.W. Peng, Y.J. Zhang, M.S. Chen, L. Xu, H.H. Liang, X.J. Lin, R.P. Guo, Y.Q. Zhang, W.Y. Lau, Radiofrequency ablation with or without transcatheter arterial chemoembolization in the treatment of hepatocellular carcinoma: A prospective randomized trial, J. Clin. Oncol. 31 (2013) 426–432. https://doi.org/10.1200/JCO.2012.42.9936.

[10] A. Liccioni, M. Reig, J. Bruix, Treatment of hepatocellular carcinoma, Dig. Dis. 32 (2014) 554–563. https://doi.org/10.1159/000360501.

[11] K. Hasegawa, T. Aoki, T. Ishizawa, J. Kaneko, Y. Sakamoto, Y. Sugawara, N. Kokudo, Comparison of the therapeutic outcomes between surgical resection and percutaneous ablation for small hepatocellular carcinoma., Ann. Surg. Oncol. 21 Suppl 3 (2014) S348–55. https://doi.org/10.1245/s10434-014-3585-x.

[12] J.C. Nault, O. Sutter, P. Nahon, N. Ganne-Carrié, O. Séror, Percutaneous treatment of hepatocellular carcinoma: State of the art and innovations, J. Hepatol. 68 (2018) 783–797. https://doi.org/10.1016/j.jhep.2017.10.004.

[13] C. Yuan, Z. Wang, D. Gu, J. Tian, P. Zhao, J. Wei, X. Yang, X. Hao, D. Dong, N. He, Y. Sun, W. Gao, J. Feng, Prediction early recurrence of hepatocellular carcinoma eligible for curative ablation using a Radiomics nomogram, Cancer Imaging. 19 (2019). https://doi.org/10.1186/s40644-019-0207-7.

[14] N. Portolani, A. Coniglio, S. Ghidoni, M. Giovanelli, A. Benetti, G.A.M. Tiberio, S.M. Giulini, Early and late recurrence after liver resection for hepatocellular carcinoma: Prognostic and therapeutic implications, Ann. Surg. 243 (2006) 229–235. https://doi.org/10.1097/01.sla.0000197706.21803.a1.

[15] J.C. Wu, Y.H. Huang, G.Y. Chau, C.W. Su, C.R. Lai, P.C. Lee, T.I. Huo, I.J. Sheen, S.D. Lee, W.Y. Lui, Risk factors for early and late recurrence in hepatitis B-related hepatocellular carcinoma, J. Hepatol. 51 (2009) 890–897. https://doi.org/10.1016/j.jhep.2009.07.009.

[16] A.W.H. Chan, J. Zhong, S. Berhane, H. Toyoda, A. Cucchetti, K.Q. Shi, T. Tada, C.C.N. Chong, B. De Xiang, L.Q. Li, P.B.S. Lai, V. Mazzaferro, M. García-Fiñana, M. Kudo, T. Kumada, S. Roayaie, P.J. Johnson, Development of pre and post-operative models to predict early recurrence of hepatocellular carcinoma after surgical resection, J. Hepatol. 69 (2018) 1284–1293. https://doi.org/10.1016/j.jhep.2018.08.027.

[17] S.F. Ang, E.S.H. Ng, H. Li, Y.H. Ong, S.P. Choo, J. Ngeow, H.C. Toh, K.H. Lim, H.Y. Yap, C.K. Tan, L.L.P.J. Ooi, A.Y.F. Chung, P.K.H. Chow, K.F. Foo, M.H. Tan, P.C. Cheow, The Singapore Liver Cancer Recurrence (SLICER) Score for relapse prediction in patients with surgically resected hepatocellular carcinoma, PLoS One. 10 (2015). https://doi.org/10.1371/journal.pone.0118658.

[18] J.H. Shim, M.J. Jun, S. Han, Y.J. Lee, S.G. Lee, K.M. Kim, Y.S. Lim, H.C. Lee, Prognostic Nomograms for Prediction of Recurrence and Survival after Curative Liver Resection for Hepatocellular Carcinoma, Ann. Surg. 261 (2015) 939–946. https://doi.org/10.1097/SLA.0000000000000747.

[19] S. Huang, G.Q. Huang, G.Q. Zhu, W.Y. Liu, J. You, K.Q. Shi, X.B. Wang, H.Y. Che, G.L. Chen, J.F. Fang, Y. Zhou, M.T. Zhou, Y.P. Chen, M. Braddock, M.H. Zheng, Establishment and validation of SSCLIP scoring system to estimate survival in hepatocellular carcinoma patients who received curative liver resection, PLoS One. 10 (2015). https://doi.org/10.1371/journal.pone.0129000.

[20] J.F. Qiu, J.Z. Ye, X.Z. Feng, Y.P. Qi, L. Ma, W.P. Yuan, J.H. Zhong, Z.M. zhang, B. De Xiang, L.Q. Li, Pre- and post-operative HBsAg levels may predict recurrence and survival after curative resection in patients with HBV-associated hepatocellular carcinoma, J. Surg. Oncol. 116 (2017) 140–148. https://doi.org/10.1002/jso.24628.

[21] W. He, B. Peng, Y. Tang, J. Yang, Y. Zheng, J. Qiu, R. Zou, J. Shen, B. Li, Y. Yuan, Nomogram to Predict Survival of Patients With Recurrence of Hepatocellular Carcinoma After Surgery, Clin. Gastroenterol. Hepatol. 16 (2018) 756–764.e10. https://doi.org/10.1016/j.cgh.2017.12.002.

[22] S.C. Lee, H.T. Tan, M.C.M. Chung, Prognostic biomarkers for prediction of recurrence of hepatocellular carcinoma: Current status and future prospects, World J. Gastroenterol. 20 (2014) 3112–3124. https://doi.org/10.3748/wjg.v20.i12.3112.

[23] Y.X. Pan, J.C. Chen, A.P. Fang, X.H. Wang, J. Bin Chen, J.C. Wang, W. He, Y.Z. Fu, L. Xu, M.S. Chen, Y.J. Zhang, Q.J. Li, Z.G. Zhou, A nomogram predicting the recurrence of hepatocellular carcinoma in patients after laparoscopic hepatectomy, Cancer Commun. 39 (2019) 55. https://doi.org/10.1186/s40880-019-0404-6.

[24] T. Barrett, D.B. Troup, S.E. Wilhite, P. Ledoux, D. Rudnev, C. Evangelista, I.F. Kim, A. Soboleva, M. Tomashevsky, K.A. Marshall, K.H. Phillippy, P.M. Sherman, R.N. Muertter, R. Edgar, NCBI GEO: archive for high-throughput functional genomic data., Nucleic Acids Res. 37 (2009) D885–90. https://doi.org/10.1093/nar/gkn764.

[25] K. Tomczak, P. Czerwińska, M. Wiznerowicz, The Cancer Genome Atlas (TCGA): An immeasurable source of knowledge, Wspolczesna Onkol. 1A (2015) A68–A77. https://doi.org/10.5114/wo.2014.47136.

[26] Z. Liu, Y. Wang, C. Dou, L. Sun, Q. Li, L. Wang, Q. Xu, W. Yang, Q. Liu, K. Tu, MicroRNA-1468 promotes tumor progression by activating PPAR-γ-mediated AKT signaling in human hepatocellular carcinoma., J. Exp. Clin. Cancer Res. 37 (2018) 49. https://doi.org/10.1186/s13046-018-0717-3.

[27] J. Wang, X. Chen, Y. Tian, G. Zhu, Y. Qin, X. Chen, L. Pi, M. Wei, G. Liu, Z. Li, C. Chen, Y. Lv, G. Cai, Six-gene signature for predicting survival in patients with head and neck squamous cell carcinoma, Aging (Albany. NY). 12 (2020) 767–783. https://doi.org/10.18632/aging.102655.

[28] Y. Ding, J.L. Yan, A.N. Fang, W.F. Zhou, L. Huang, Circulating miRNAs as novel diagnostic biomarkers in hepatocellular carcinoma detection: A meta-analysis based on 24 articles, Oncotarget. 8 (2017) 66402–66413. https://doi.org/10.18632/oncotarget.18949.

[29] F. Liu, H. Li, H. Chang, J. Wang, J. Lu, Identification of hepatocellular carcinoma-associated hub genes and pathways by integrated microarray analysis., Tumori. 101 (2015) 206–14. https://doi.org/10.5301/tj.5000241.

[30] J. Xu, C. Wu, X. Che, L. Wang, D. Yu, T. Zhang, L. Huang, H. Li, W. Tan, C. Wang, D. Lin, Circulating MicroRNAs, miR-21, miR-122, and miR-223, in patients with hepatocellular carcinoma or chronic hepatitis, Mol. Carcinog. 50 (2011) 136–142. https://doi.org/10.1002/mc.20712.

[31] Y. Zhao, F. Li, X. Zhang, A. Liu, J. Qi, H. Cui, P. Zhao, MicroRNA-194 acts as a prognostic marker and inhibits proliferation in hepatocellular carcinoma by targeting MAP4K4., Int. J. Clin. Exp. Pathol. 8 (2015) 12446–54. http://www.ncbi.nlm.nih.gov/pubmed/26722431 (accessed June 6, 2020).

[32] H.Y. Lim, I. Sohn, S. Deng, J. Lee, S.H. Jung, M. Mao, J. Xu, K. Wang, S. Shi, J.W. Joh, Y. La Choi, C.K. Park, Prediction of disease-free survival in hepatocellular carcinoma by gene expression profiling, Ann. Surg. Oncol. 20 (2013) 3747–3753. https://doi.org/10.1245/s10434-013-3070-y.

[33] M.K. Bhasin, K. Ndebele, O. Bucur, E.U. Yee, H.H. Otu, J. Plati, A. Bullock, X. Gu, E. Castan, P. Zhang, R. Najarian, M.S. Muraru, R. Miksad, R. Khosravi-Far, T.A. Libermann, Meta-analysis of transcriptome data identifies a novel 5-gene pancreatic adenocarcinoma classifier, Oncotarget. 7 (2016) 23263–23281. https://doi.org/10.18632/oncotarget.8139.

[34] R.-M. Chang, S. Xiao, X. Lei, H. Yang, F. Fang, L.-Y. Yang, miRNA-487a Promotes Proliferation and Metastasis in Hepatocellular Carcinoma., Clin. Cancer Res. 23 (2017) 2593–2604. https://doi.org/10.1158/1078-0432.CCR-16-0851.

[35] H.L. Jia, Q.H. Ye, L.X. Qin, A. Budhu, M. Forgues, Y. Chen, Y.K. Liu, H.C. Sun, L. Wang, H.Z. Lu, F. Shen, Z.Y. Tang, W.W. Xin, Gene expression profiling reveals potential biomarkers of human hepatocellular carcinoma, Clin. Cancer Res. 13 (2007) 1133–1139. https://doi.org/10.1158/1078-0432.CCR-06-1025.

[36] Z. Makowska, T. Boldanova, D. Adametz, L. Quagliata, J.E. Vogt, M.T. Dill, M.S. Matter, V. Roth, L. Terracciano, M.H. Heim, Gene expression analysis of biopsy samples reveals critical limitations of transcriptome-based molecular classifications of hepatocellular carcinoma, J. Pathol. Clin. Res. 2 (2016) 80–92. https://doi.org/10.1002/cjp2.37.

[37] Y. Zheng, Y. Liu, S. Zhao, Z. Zheng, C. Shen, L. An, Y. Yuan, Large-scale analysis reveals a novel risk score to predict overall survival in hepatocellular carcinoma, Cancer Manag. Res. 10 (2018) 6079–6096. https://doi.org/10.2147/CMAR.S181396.

[38] B. Gao, S. Ning, J. Li, H. Liu, W. Wei, F. Wu, Y. Tang, Y. Feng, K. Li, L. Zhang, Integrated analysis of differentially expressed mRNAs and miRNAs between hepatocellular carcinoma and their matched adjacent normal liver tissues., Oncol. Rep. 34 (2015) 325–33. https://doi.org/10.3892/or.2015.3968.

[39] R.-T. Xie, X.-L. Cong, X.-M. Zhong, P. Luo, H.-Q. Yang, G.-X. Lu, P. Luo, Z.-Y. Chang, R. Sun, T.-M. Wu, Z.-W. Lv, D. Fu, Y.-S. Ma, MicroRNA-33a downregulation is associated with tumorigenesis and poor prognosis in patients with hepatocellular carcinoma., Oncol. Lett. 15 (2018) 4571–4577. https://doi.org/10.3892/ol.2018.7892.

[40] G. Li, Q. Shen, C. Li, D. Li, J. Chen, M. He, Identification of circulating MicroRNAs as novel potential biomarkers for hepatocellular carcinoma detection: a systematic review and meta-analysis, Clin. Transl. Oncol. 17 (2015) 684–693. https://doi.org/10.1007/s12094-015-1294-y.

[41] D. Chakroborty, M.R. Emani, R. Klén, C. Böckelman, J. Hagström, C. Haglund, A. Ristimäki, R. Lahesmaa, L.L. Elo, L1TD1 - A prognostic marker for colon cancer, BMC Cancer. 19 (2019) 727. https://doi.org/10.1186/s12885-019-5952-2.

[42] D.G.P. van IJzendoorn, K. Szuhai, I.H. Briaire-De Bruijn, M. Kostine, M.L. Kuijjer, J.V.M.G. Bovée, Machine learning analysis of gene expression data reveals novel diagnostic and prognostic biomarkers and identifies therapeutic targets for soft tissue sarcomas, PLoS Comput. Biol. 15 (2019). https://doi.org/10.1371/journal.pcbi.1006826.

[43] J. Feng, R. Zhu, C. Chang, L. Yu, F. Cao, G. Zhu, F. Chen, H. Xia, F. Lv, S. Zhang, L. Sun, CK19 and glypican 3 expression profiling in the prognostic indication for patients with HCC after surgical resection, PLoS One. 11 (2016). https://doi.org/10.1371/journal.pone.0151501.

[44] F. Chen, X.-F. Li, D.-S. Fu, J.-G. Huang, S.-E. Yang, Clinical potential of miRNA-221 as a novel prognostic biomarker for hepatocellular carcinoma., Cancer Biomark. 18 (2017) 209–214. https://doi.org/10.3233/CBM-161671.

[45] S. Bhalla, H. Kaur, R. Kaur, S. Sharma, G.P.S. Raghava, Expression based biomarkers and models to classify early and late-stage samples of Papillary Thyroid Carcinoma, PLoS One. 15 (2020) e0231629. https://doi.org/10.1371/journal.pone.0231629.

[46] J. Ji, H. Chen, X.P. Liu, Y.H. Wang, C.L. Luo, W.W. Zhang, W. Xie, F.B. Wang, A miRNA combination as promising biomarker for hepatocellular carcinoma diagnosis: A study based on bioinformatics analysis, J. Cancer. 9 (2018) 3435–3446. https://doi.org/10.7150/jca.26101.

[47] X. Xu, Y. Zhang, L. Zou, M. Wang, A. Li, A gene signature for breast cancer prognosis using support vector machine, in: 2012 5th Int. Conf. Biomed. Eng. Informatics, BMEI 2012, 2012: pp. 928–931. https://doi.org/10.1109/BMEI.2012.6513032.

[48] J. Kong, T. Wang, S. Shen, Z. Zhang, X. Yang, W. Wang, A genomic-clinical nomogram predicting recurrence-free survival for patients diagnosed with hepatocellular carcinoma, PeerJ. 2019 (2019). https://doi.org/10.7717/peerj.7942.

[49] Y. Hasin, M. Seldin, A. Lusis, Multi-omics approaches to disease, Genome Biol. 18 (2017) 1–15. https://doi.org/10.1186/s13059-017-1215-1.

[50] G. de Anda-Jáuregui, E. Hernández-Lemus, Computational Oncology in the Multi-Omics Era: State of the Art, Front. Oncol. 10 (2020) 423. https://doi.org/10.3389/fonc.2020.00423.

[51] C. S, H. MI, A. M, S. HU, Onco-Multi-OMICS Approach: A New Frontier in Cancer Research, Biomed Res. Int. 2018 (2018). https://doi.org/10.1155/2018/9836256.

[52] H. Kaur, S. Bhalla, G.P.S. Raghava, Classification of early and late stage liver hepatocellular carcinoma patients from their genomics and epigenomics profiles., PLoS One. 14 (2019) e0221476. https://doi.org/10.1371/journal.pone.0221476.

[53] M.A. Jensen, V. Ferretti, R.L. Grossman, L.M. Staudt, The NCI Genomic Data Commons as an engine for precision medicine, Blood. 130 (2017) 453–459. https://doi.org/10.1182/blood-2017-03-735654.

[54] S. Roessler, H.L. Jia, A. Budhu, M. Forgues, Q.H. Ye, J.S. Lee, S.S. Thorgeirsson, Z. Sun, Z.Y. Tang, L.X. Qin, X.W. Wang, A unique metastasis gene signature enables prediction of tumor relapse in early-stage hepatocellular carcinoma patients, Cancer Res. 70 (2010) 10202–10212. https://doi.org/10.1158/0008-5472.CAN-10-2607.

[55] S. Roessler, E.L. Long, A. Budhu, Y. Chen, X. Zhao, J. Ji, R. Walker, H.L. Jia, Q.H. Ye, L.X. Qin, Z.Y. Tang, P. He, K.W. Hunter, S.S. Thorgeirsson, P.S. Meltzer, X.W. Wang, Integrative genomic identification of genes on 8p associated with hepatocellular carcinoma progression and patient survival, Gastroenterology. 142 (2012). https://doi.org/10.1053/j.gastro.2011.12.039.

[56] A. Budhu, H.L. Jia, M. Forgues, C.G. Liu, D. Goldstein, A. Lam, K.A. Zanetti, Q.H. Ye, L.X. Qin, C.M. Croce, Z.Y. Tang, W.W. Xin, Identification of metastasis-related microRNAs in hepatocellular carcinoma, Hepatology. 47 (2008) 897–907. https://doi.org/10.1002/hep.22160.

[57] A. Budhu, S. Roessler, X. Zhao, Z. Yu, M. Forgues, J. Ji, E. Karoly, L.X. Qin, Q.H. Ye, H.L. Jia, J. Fan, H.C. Sun, Z.Y. Tang, X.W. Wang, Integrated metabolite and gene expression profiles identify lipid biomarkers associated with progression of hepatocellular carcinoma and patient outcomes, Gastroenterology. 144 (2013). https://doi.org/10.1053/j.gastro.2013.01.054.

[58] S. Davis, Using the GEOquery package, 2013. http://www.ncbi.nih.gov/geo (accessed June 15, 2020).

[59] S. Davis, P.S. Meltzer, GEOquery: a bridge between the Gene Expression Omnibus (GEO) and BioConductor, Academic.Oup.Com. 23 (2007) 1846–1847. https://doi.org/10.1093/bioinformatics/btm254.

[60] B.S. Carvalho, R.A. Irizarry, A framework for oligonucleotide microarray preprocessing., Bioinformatics. 26 (2010) 2363–7. https://doi.org/10.1093/bioinformatics/btq431.

[61] B.S. Carvalho, Working with Oligonucleotide Arrays., Methods Mol. Biol. 1418 (2016) 145–59. https://doi.org/10.1007/978-1-4939-3578-9_7.

[62] H.-C. Huang, L.-X. Qin, Empirical evaluation of data normalization methods for molecular classification., PeerJ. 6 (2018) e4584. https://doi.org/10.7717/peerj.4584.

[63] C.B. Pedersen, F.C. Nielsen, M. Rossing, L.R. Olsen, Using microarray-based subtyping methods for breast cancer in the era of high-throughput RNA sequencing, Mol. Oncol. 12 (2018) 2136–2146. https://doi.org/10.1002/1878-0261.12389.

[64] M. Kuhn, The caret Package, 2011. http://citeseerx.ist.psu.edu/viewdoc/download?doi=10.1.1.216.2142&rep=rep1&type=pdf (accessed June 10, 2020).

[65] T. Therneau, A package for survival analysis in S. R package version 2.37–7, (2014).

[66] A. Kassambara, M. Kosinski, P.B.-R. package version 0.3, undefined 2017, survminer: Drawing Survival Curves using’ggplot2’, (n.d.).

[67] S. Bhalla, H. Kaur, A. Dhall, G.P.S. Raghava, Prediction and Analysis of Skin Cancer Progression using Genomics Profiles of Patients, Sci. Rep. 9 (2019) 1–16. https://doi.org/10.1038/s41598-019-52134-4.

[68] F. Pedregosa FABIANPEDREGOSA, V. Michel, O. Grisel OLIVIERGRISEL, M. Blondel, P. Prettenhofer, R. Weiss, J. Vanderplas, D. Cournapeau, F. Pedregosa, G. Varoquaux, A. Gramfort, B. Thirion, O. Grisel, V. Dubourg, A. Passos, M. Brucher, M. Perrot and Édouardand, and Édouard Duchesnay, Fré. Duchesnay EDOUARDDUCHESNAY, Scikit-learn: Machine Learning in Python Gaёl Varoquaux Bertrand Thirion Vincent Dubourg Alexandre Passos PEDREGOSA, VAROQUAUX, GRAMFORT ET AL. Matthieu Perrot, 2011. http://scikit-learn.sourceforge.net. (accessed June 9, 2020).

[69] A. Gulli, S. Pal, Deep learning with Keras, 2017. https://books.google.com/books?hl+en&lr=&id=20EwDwAAQBAJ&oi=fnd&pg=PP1&dq=keras+python+package&ots=lHhwcn8UR1&sig=YQ_Yh9llInkQUGWEYy1tMYOBp8s (accessed June 11, 2020).

[70] L. Zhang, C. Lv, Y. Jin, G. Cheng, Y. Fu, D. Yuan, Y. Tao, Y. Guo, X. Ni, T. Shi, Deep Learning-Based Multi-Omics Data Integration Reveals Two Prognostic Subtypes in High-Risk Neuroblastoma, Front. Genet. 9 (2018) 477. https://doi.org/10.3389/fgene.2018.00477.

[71] K. Chaudhary, O.B. Poirion, L. Lu, L.X. Garmire, Deep learning–based multi-omics integration robustly predicts survival in liver cancer, Clin. Cancer Res. 24 (2018) 1248–1259. https://doi.org/10.1158/1078-0432.CCR-17-0853.

[72] C. Sticht, C. De La Torre, A. Parveen, N. Gretz, Mirwalk: An online resource for prediction of microrna binding sites, PLoS One. 13 (2018). https://doi.org/10.1371/journal.pone.0206239.

[73] H. Dweep, N. Gretz, C. Sticht, MiRWalk database for miRNA-target interactions, Methods Mol. Biol. 1182 (2014) 289–305. https://doi.org/10.1007/978-1-4939-1062-5_25.

[74] G. Stelzer, N. Rosen, I. Plaschkes, S. Zimmerman, M. Twik, S. Fishilevich, T.I. Stein, R. Nudel, I. Lieder, Y. Mazor, S. Kaplan, D. Dahary, D. Warshawsky, Y. Guan□Golan, A. Kohn, N. Rappaport, M. Safran, D. Lancet, The GeneCards Suite: From Gene Data Mining to Disease Genome Sequence Analyses, Curr. Protoc. Bioinforma. 54 (2016) 1.30.1–1.30.33. https://doi.org/10.1002/cpbi.5.

[75] L.L. Studach, S. Menne, S. Cairo, M.A. Buendia, R.L. Hullinger, L. Lefrançois, P. Merle, O.M. Andrisani, Subset of Suz12/PRC2 target genes is activated during hepatitis B virus replication and liver carcinogenesis associated with HBV X protein, Hepatology. 56 (2012) 1240–1251. https://doi.org/10.1002/hep.25781.

[76] C. Xue, K. Wang, X. Jiang, C. Gu, G. Yu, Y. Zhong, S. Liu, Y. Nie, Y. Zhou, H. Yang, The down-regulation of SUZ12 accelerates the migration and invasion of liver cancer cells via activating ERK1/2 pathway, J. Cancer. 10 (2019) 1375–1384. https://doi.org/10.7150/jca.29932.

[77] J. Zhou, H.C. Sun, Z. Wang, W.M. Cong, J.H. Wang, M.S. Zeng, J.M. Yang, P. Bie, L.X. Liu, T.F. Wen, G.H. Han, M.Q. Wang, R.B. Liu, L.G. Lu, Z.G. Ren, M.S. Chen, Z.C. Zeng, P. Liang, C.H. Liang, M. Chen, F.H. Yan, W.P. Wang, Y. Ji, W.W. Cheng, C.L. Dai, W.D. Jia, Y.M. Li, Y.X. Li, J. Liang, T.S. Liu, G.Y. Lv, Y.L. Mao, W.X. Ren, H.C. Shi, W.T. Wang, X.Y. Wang, B.C. Xing, J.M. Xu, J.Y. Yang, Y.F. Yang, S. L. Ye, Z.Y. Yin, B.H. Zhang, S.J. Zhang, W.P. Zhou, J.Y. Zhu, R. Liu, Y.H. Shi, Y.S. Xiao, Z. Dai, G.J. Teng, J.Q. Cai, W.L. Wang, J.H. Dong, Q. Li, F. Shen, S.K. Qin, J. Fan, Guidelines for diagnosis and treatment of primary liver cancer in China (2017 Edition), Liver Cancer. 7 (2018) 235–260. https://doi.org/10.1159/000488035.

[78] J. Bruix, M. Sherman, Management of hepatocellular carcinoma: An update, Hepatology. 53 (2011) 1020–1022. https://doi.org/10.1002/hep.24199.

[79] J.M. Llovet, M. Ducreux, R. Lencioni, A.M. Di Bisceglie, P.R. Galle, J.F. Dufour, T.F. Greten, E. Raymond, T. Roskams, T. De Baere, M. Ducreux, V. Mazzaferro, M. Bernardi, J. Bruix, M. Colombo, A. Zhu, EASL-EORTC Clinical Practice Guidelines: Management of hepatocellular carcinoma, J. Hepatol. 56 (2012) 908–943. https://doi.org/10.1016/j.jhep.2011.12.001.

[80] N.A. Filgueira, Hepatocellular carcinoma recurrence after liver transplantation: Risk factors, screening and clinical presentation, World J. Hepatol. 11 (2019) 261–272. https://doi.org/10.4254/wjh.v11.i3.261.

[81] H. Kaur, A. Dhall, R. Kumar, G.P.S. Raghava, Identification of Platform-Independent Diagnostic Biomarker Panel for Hepatocellular Carcinoma Using Large-Scale Transcriptomics Data, Front. Genet. 10 (2020). https://doi.org/10.3389/fgene.2019.01306.

[82] H. Kaur, S. Bhalla, D. Kaur, G.P. Raghava, CancerLivER: a database of liver cancer gene expression resources and biomarkers, Database. 2020 (2020) 12. https://doi.org/10.1093/database/baaa012.

[83] H. Kaur, S. Bhalla, D. Garg, N. Mehta, G.P.S. Raghava, Analysis and prediction of cholangiocarcinoma from transcriptomic profile of patients, J. Hepatol. 73 (2020) S16–S17. https://doi.org/10.1016/s0168-8278(20)30593-6.

[84] R. Kumar, P. Bhanti, A. Marwal, R.K. Gaur, Gene Expression-Based Supervised Classification Models for Discriminating Early- and Late-Stage Prostate Cancer, Proc. Natl. Acad. Sci. India Sect. B - Biol. Sci. (2019) 1–25. https://doi.org/10.1007/s40011-019-01127-4.

[85] S. Bhalla, K. Chaudhary, R. Kumar, M. Sehgal, H. Kaur, S. Sharma, G.P.S. Raghava, Gene expression-based biomarkers for discriminating early and late stage of clear cell renal cancer., Sci. Rep. 7 (2017) 44997. https://doi.org/10.1038/srep44997.

[86] L.L. Studach, S. Menne, S. Cairo, M.A. Buendia, R.L. Hullinger, L. Lefrançois, P. Merle, O.M. Andrisani, Subset of Suz12/PRC2 target genes is activated during hepatitis B virus replication and liver carcinogenesis associated with HBV X protein, Hepatology. 56 (2012) 1240–1251. https://doi.org/10.1002/hep.25781.

[87] S. Nakagawa, H. Okabe, Y. Sakamoto, H. Hayashi, D. Hashimoto, N. Yokoyama, K. Sakamoto, H. Kuroki, K. Mima, H. Nitta, K. Imai, A. Chikamoto, M. Watanabe, T. Beppu, H. Baba, Enhancer of Zeste Homolog 2 (EZH2) promotes progression of cholangiocarcinoma cells by regulating cell cycle and apoptosis, Ann. Surg. Oncol. 20 (2013). https://doi.org/10.1245/s10434-013-3135-y.

[88] D. Martin-Perez, E. Sanchez, L. Maestre, M.A. Piris, M. Sanchez-Beato, SUZ12 overexpression in human tumors is associated with tumoral transformation and its depletion affects Mantle cell lymphoma cell lines viability., Cancer Res. 67 (2007).

[89] Y. Wu, H. Hu, W. Zhang, Z. Li, P. Diao, D. Wang, W. Zhang, Y. Wang, J. Yang, J. Cheng, SUZ12 is a novel putative oncogene promoting tumorigenesis in head and neck squamous cell carcinoma, J. Cell. Mol. Med. 22 (2018) 3582–3594. https://doi.org/10.1111/jcmm.13638.

