## Supplementary File 2 for "Integrative multi-omics approach for stratification of tumor recurrence risk groups of Hepatocellular Carcinoma patients"

**Integrative multi-omics approach for stratification of tumor recurrence risk groups of Hepatocellular Carcinoma patients**

**Harpreet Kaur^1^, Anjali Lathwal^2^, Gajendra P.S. Raghava^2*^**

^1^	Bioinformatics Center, CSIR-Institute of Microbial Technology, Chandigarh, India

^2^	Department of Computational Biology, Indraprastha Institute of Information Technology, New Delhi, India

**Emails of Authors:**

Harpreet Kaur:

Anjali Lathwal:

Gajendra P.S. Raghava:

***	Correspondence**

Professor, Department of Computational Biology

Indraprastha Institute of Information Technology,

Okhla Industrial Estate, Phase III, New Delhi 110020,

**Supplementary Figures:**

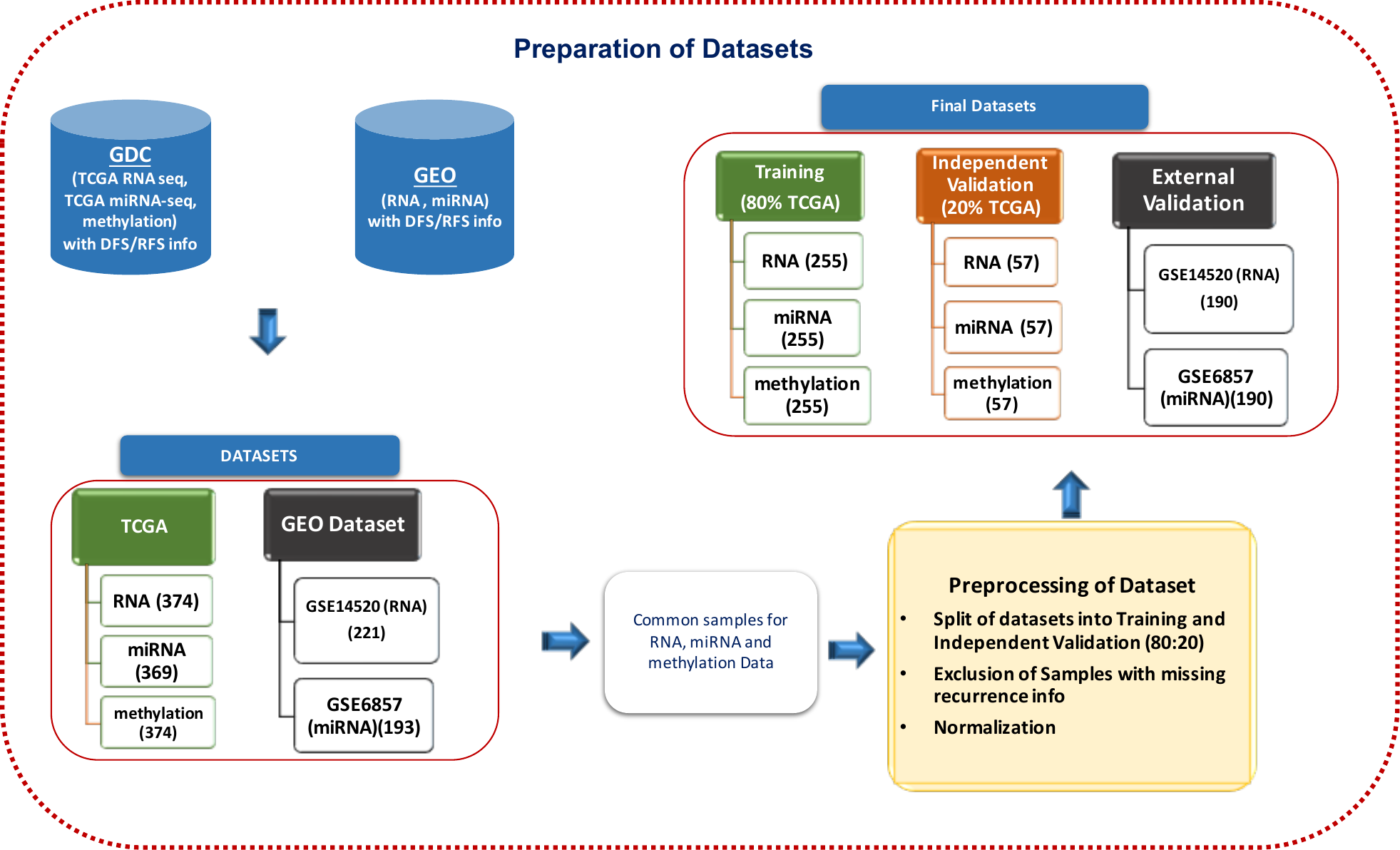

**Figure S1: Compilation of Datasets (TCGA, GEO) from the GDC and GEO Resources; the creation of training, independent and external validation datasets.**

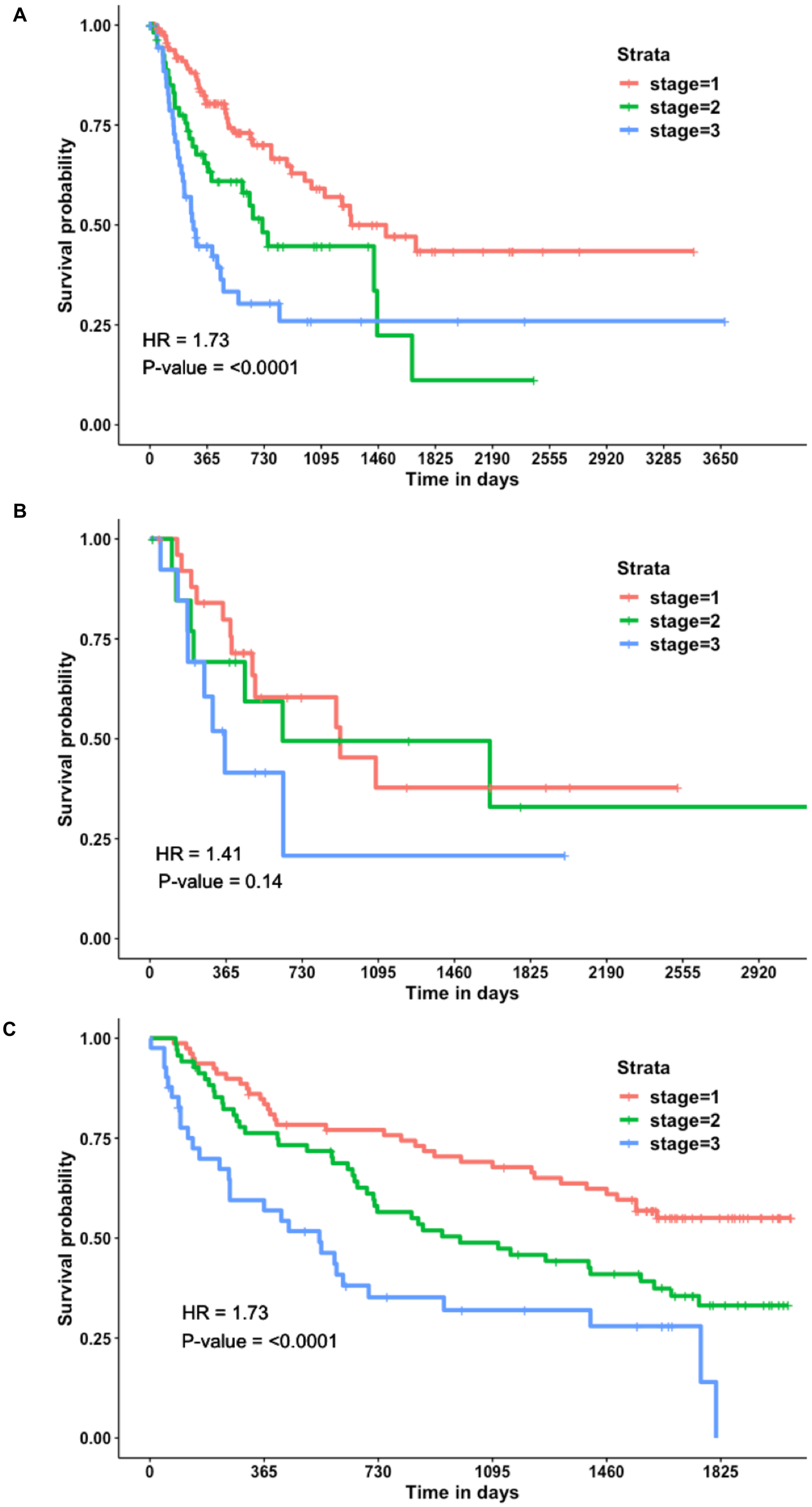

**Figure S2: Kaplan-Meier plots depict the stratification of recurrence risk groups of HCC samples based on tumor stage: (A) Training, (B) Independent, and (C) External validation dataset.**

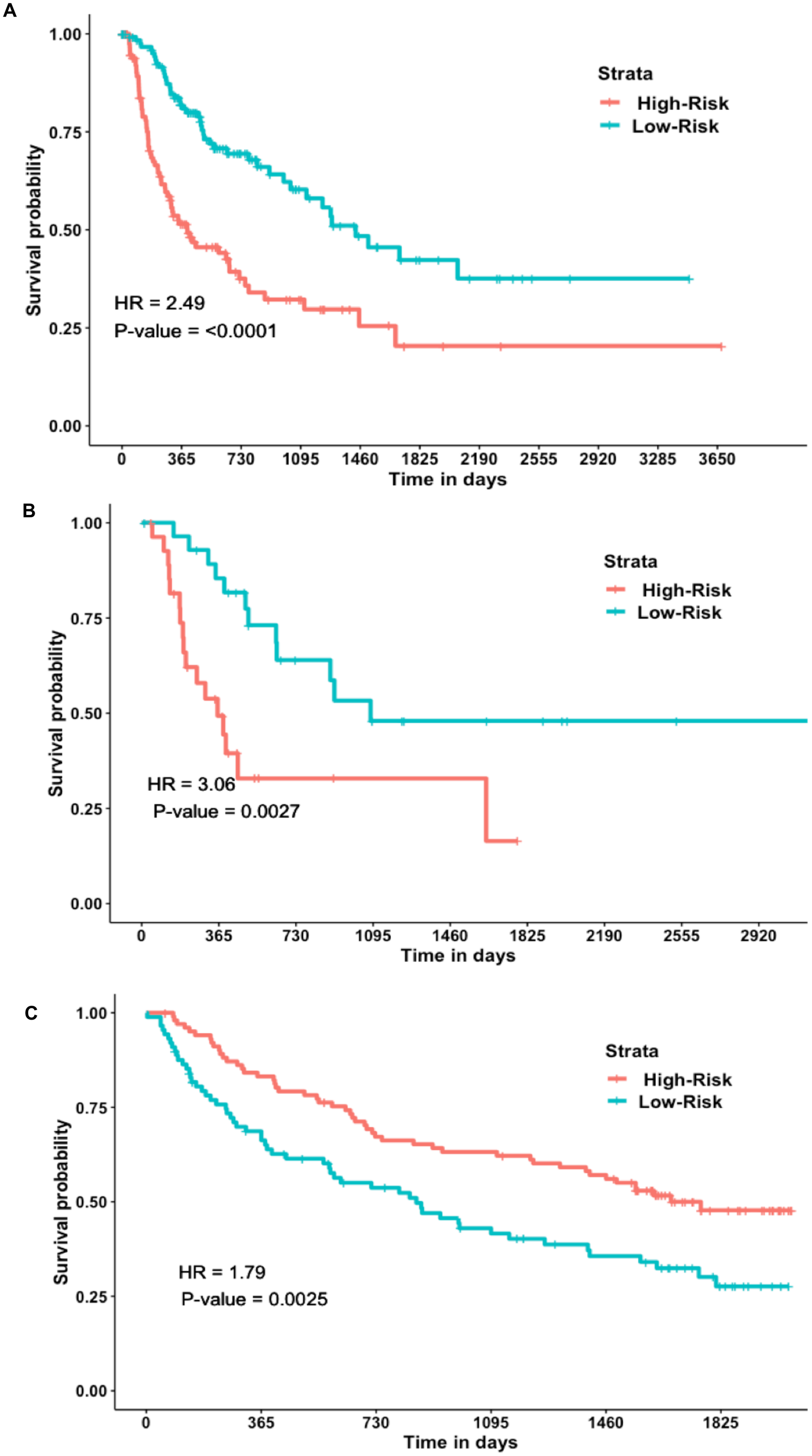

**Figure S3: Kaplan-Meier plots depict the stratification of recurrence risk groups based on the prediction with K-mean clustering based on 28 PCA features: (A) Training, (B) Independent, and (C) External validation dataset.**
